# Rhythmic light stimulation elicits multiple, concurrent neural responses that separably shape human perception

**DOI:** 10.1101/2025.11.05.686709

**Authors:** Benjamin J. Griffiths, Katharina Duecker, Camille Fakche, Laura Dugué, Andrew Quinn, Ole Jensen

## Abstract

Rhythmic light stimulation offers solutions to innumerable cognitive and neurological disorders. However, like any neuromodulatory technique, responses to rhythmic light stimulation are highly variable, producing challenges in replicating lab-based studies and translating findings to the clinic. Across three MEG/EEG experiments, we show that this variability can, in part, be attributed to rhythmic light stimulation eliciting multiple, coexisting neural responses which have separable impacts on cognition. Specifically, we find that rhythmic light stimulation produces distinct neural responses at the fundamental (*f*) and second harmonic (*2f*) frequencies, and that these responses are differentially shaped by endogenous oscillatory dynamics that vary across participants. Importantly, these responses separably contribute to perception, with harmonic gamma-band responses supporting the representation of stimulus-specific information, and the phase of harmonic alpha-band responses causally contributing to near-threshold visual perception. We reproduce these effects across datasets, paradigms, and oscillatory bands, suggesting that the multiplex oscillatory responses elicited by rhythmic light stimulation are a robust and pervasive phenomenon. We propose that the complexity of neural responses to rhythmic stimulation can explain why there is substantial variability between studies using these techniques, and that understanding these complex responses may help advance neuromodulatory technologies for both fundamental and clinical neuroscience.

**Supplementary Material:** All supplementary material can be found at the end of this document.

## Main Text

Rhythmic light stimulation modulates brain activity^1–4^, enabling us to identify causal mechanisms underpinning cognition^5,6^ and pioneer game-changing neurological interventions^7–9^. Compared to other rhythmic neuromodulatory approaches (e.g., transcranial magnetic stimulation [TMS]; deep brain stimulation [DBS]), rhythmic light stimulation offers high clinical throughput (i.e., number of procedures per day) at low cost, meaning it can tackle global neurological health issues at scale. To do this, however, rhythmic light stimulation needs to be accurate^10,11^ and precise^12–14^, modulating the neural target and nothing more. Given the brain’s complex, non-linear dynamics^15,16^ however, we propose that such specificity is unfeasible. Instead, more success may be found in understanding the myriad neural responses elicited by a rhythmic input and accounting for how each of these responses separably influences cognition.

Neural oscillations are a common target of rhythmic light stimulation. Neural oscillations correlate with innumerable cognitive functions^6,17,18^ (e.g., visual perception^19,20^, memory^21,22^, and decision-making^23,24^) and their disruption coincides with numerous major neurological and psychiatric disorders^25,26^ (including Alzheimer’s Disease^27,28^, Parkinson’s Disease^26,29^, depression^30,31^, and schizophrenia^32,33^). Therefore, there is much to be gained by causally controlling oscillations. While rhythmic light stimulation has been suggested to do this^6^, the outcomes of such studies are inconsistent: promising results^28,34^ are followed by failures to replicate^35,36^. It is unclear why these inconsistencies arise^37^, but an explanation may be found in the diversity of oscillatory responses induced by rhythmic light stimulation.

Oscillatory responses to rhythmic light stimulation are thought to be strongest when the stimulation frequency matches the resonant frequency of the target oscillation^1,38,39^. This principle, formalised in the concept of Arnold Tongues^40^, underpins much of the logic behind rhythmic brain stimulation^1,38,39^. However, Arnold Tongues also occur at integer multiples of the input frequency^40,41^ (see Figure 1A) – a phenomenon evident throughout biology, including in tumour-supressing proteins^42^, chick hearts^43^, the retina of salamanders^44^, and neural phase-phase coupling^45,46^. Therefore, it is plausible to suggest that oscillatory responses to rhythmic light stimulation may arise in higher harmonics as well as at the stimulation frequency, opening the door to unintentional side-effects from stimulation if these higher harmonic responses impact cognition.

**Figure 1.**
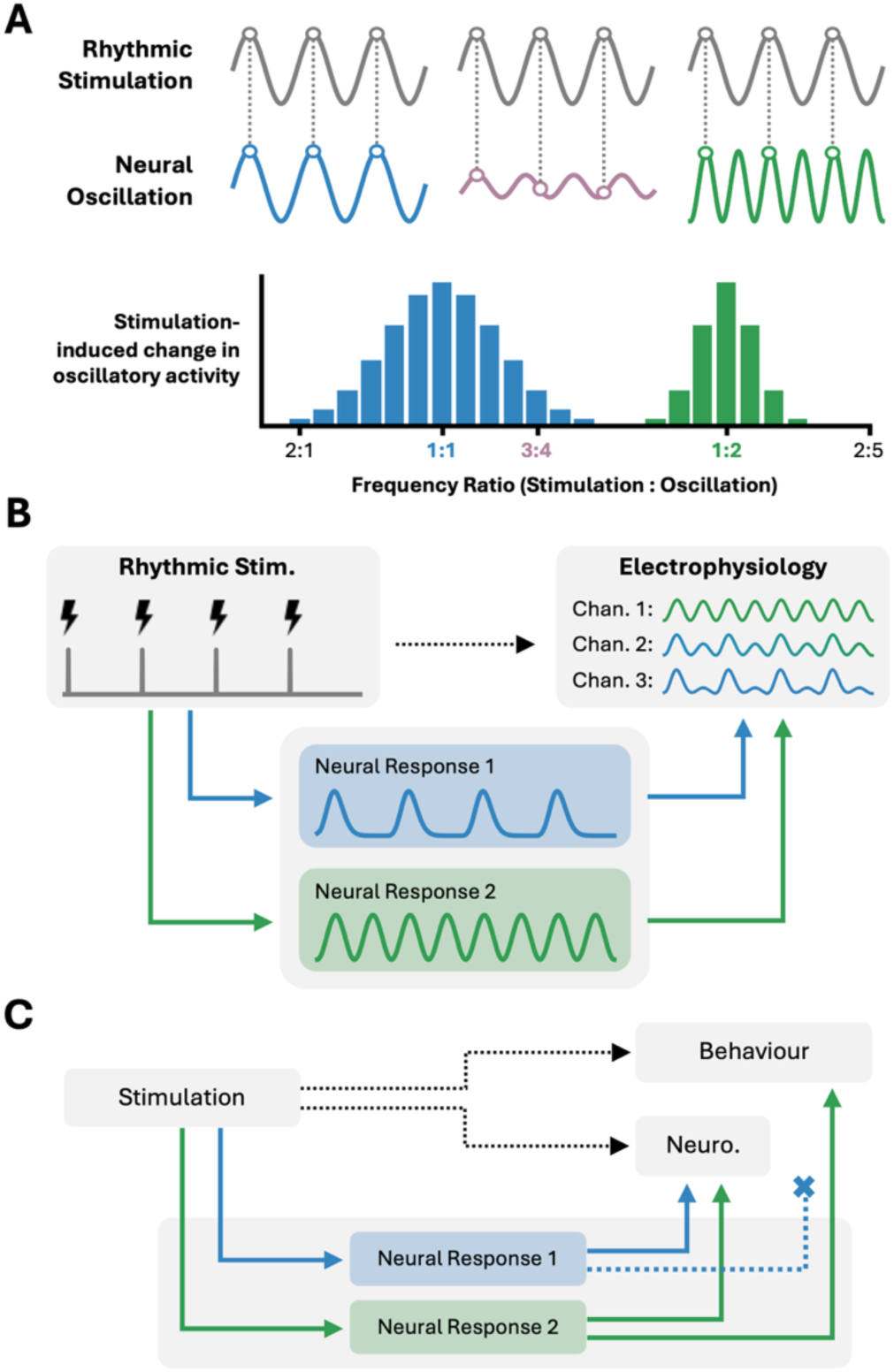
Key concepts. **(A) Harmonic responses to rhythmic stimulation**. Rhythmic stimulation modulates oscillatory activity when the frequencies approximately align (blue) or when the oscillatory frequency is an integer multiple of the stimulation frequency (green). Stimulation frequencies between these two ratios (purple) do not align with the oscillatory frequency, and therefore struggle to amplify the oscillations. **(B) Emergence of stimulation-induced complex rhythms in electrophysiology.** In this example, rhythmic stimulation produces two neural responses in neighbouring regions: one at the fundamental frequency, one at the second harmonic. The spatial proximity of the neural responses mean they sum together and present as a multiplex wave on a recorded channel. None of the raw measures reflect the true neural responses (though spatial and spectral differences between the outputs can be used to reconstruct the neural responses; not visualised). **(C) The relationship between neural and behavioural responses to stimulation.** Stimulation elicits multiple latent responses within the brain. These responses sum together to produce measurable effects in neuroimaging and behaviour, but the way in which they sum together differ between measurements. In this instance, “Neural Response 1” contributes to the neural measure, but does not affect behaviour. This means the raw output from one measure will not necessarily predict another output.

To further complicate matters, neural responses to stimulation may become multiplex if the higher harmonic responses interact or summate with other responses^4^ (e.g., stimulation-evoked responses at the fundamental^47^). Take an example in which rhythmic stimulation at 4Hz induces two responses: a steady-state visually-evoked response at the fundamental (*f*; 4Hz) and an oscillatory response at the second harmonic (2*f*; 8Hz). If the neural responses come from overlapping sources, EEG electrodes will capture a multiplex signal that reflects the weighted sum of the 4Hz and 8Hz responses (see Figure 1B). This composite signal, as captured by EEG, is not necessarily the driver of any observed change in behaviour^48^. Rather, it could be that (i) only one of the neural responses impacts behaviour (see Figure 1C), (ii) the responses have separate influences over behaviour^49–51^, or (iii) they interact non-linearly to modulate behaviour^34,52,53^. Failure to account for these possibilities will lead to erroneous interpretations of results and consequent failures to replicate. To circumvent this, we need to recover the underlying neural responses to stimulation from measured activity and relate the recovered responses to changes in cognition and/or behaviour. Here, we aim to do just that.

### Current Experiment

We hypothesise that (i) rhythmic light stimulation elicits a harmonic oscillatory response that is distinct from the well-documented response at the fundamental frequency^1,3,4,38,47,54,55^, and (ii) the fundamental and higher harmonic responses have separable influences over human perception.

Analytically, these hypotheses are challenging as traditional oscillatory analyses rely on the Fourier transform^56^, which cannot distinguish whether power increases at fundamental and harmonic frequencies reflect (i) distinct neural oscillations, or (ii) a single, non-sinusoidal oscillation^26,57–59^. This ambiguity complicates the reinterpretation of prior EEG research on harmonics. For example, while several studies^60–65^ have found spatially separable harmonic components in FFT-derived power spectra following rhythmic stimulation, this does not mean that rhythmic stimulation has induced a neural oscillation at the harmonic as non-sinusoidal responses to rhythmic stimulation also produce spatially separable harmonic patterns in FFT-power spectra (see Supp. Figure 1). To circumvent this problem, we use an alternative approach: empirical mode decomposition^66,67^ (EMD). EMD separates the recorded electrophysiological signal into distinct rhythmic components, termed intrinsic mode functions (IMFs; see Figure 2E). Critically, overlapping oscillatory responses at the fundamental and higher harmonic frequencies will be separated as distinct IMFs if the peaks of both components are visible in the summation. In contrast, singular oscillations with a non-sinusoidal waveform will present as a single rhythm^68^. The fundamental and harmonic IMFs we resolve can then be analysed separately, enabling us to identify if they operate separably on a neural and/or behavioural level. To find evidence that the IMFs are separable (e.g., spatially, temporally, or functionally) from one another would mean that multiplex neural responses to rhythmic light stimulation should not be treated as a singular entity, but rather a summation of multiple concurrent responses to stimulation.

**Figure 2.**
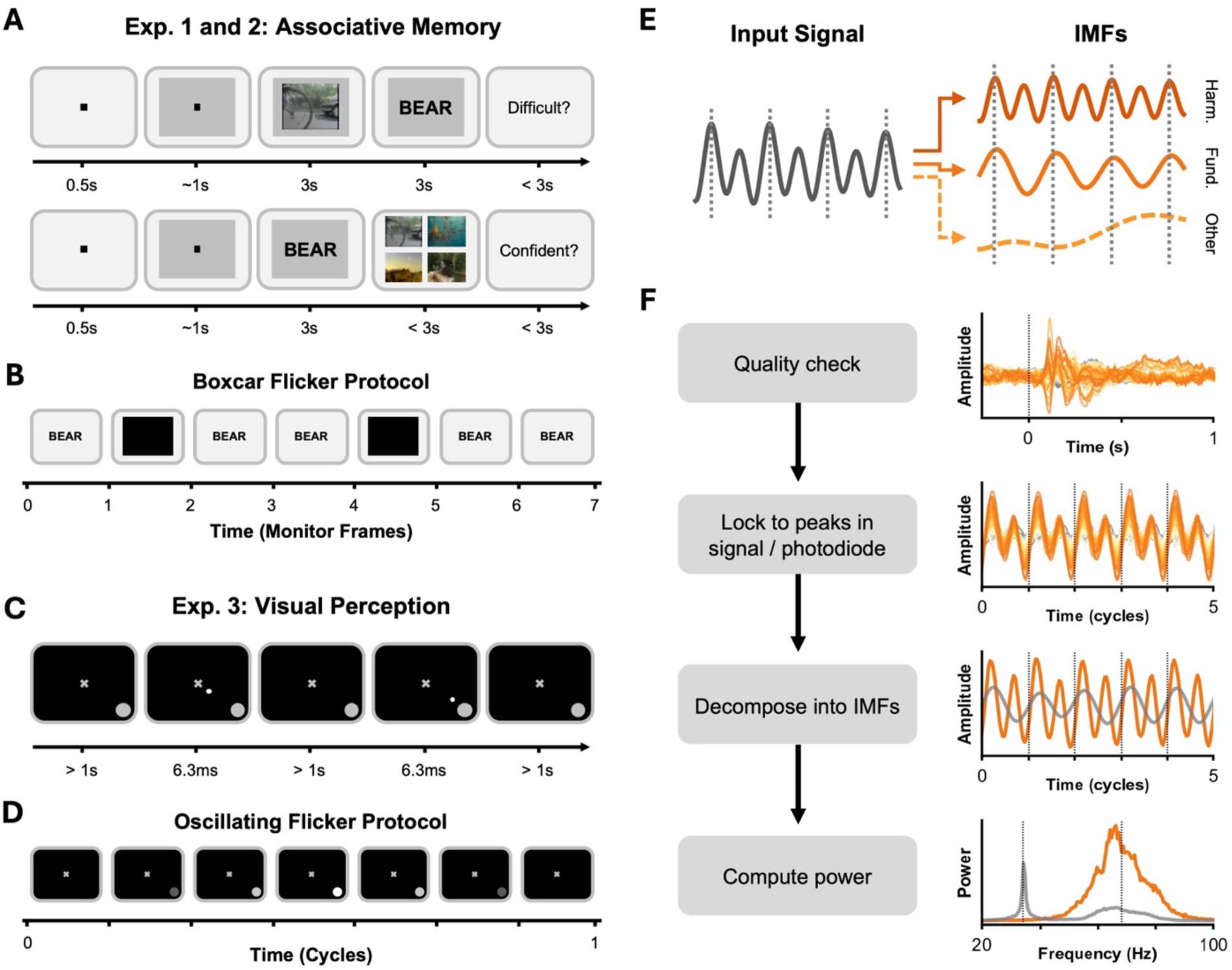
Experimental and analytical approach. **(A)** Experiments 1 and 2 used an associative memory paradigm in which participants associated between a video (one of four) and a unique word. Later, they recalled the video using the word as a cue. Full details on the task/flicker are reported in GriDiths et al. (2023; bioRxiv). **(B)** The flicker was delivered by periodically turning a patch on the screen black. The visualisation describes how this was achieved for 40Hz in Experiment 1, which used a 120Hz monitor. **(C)** Experiment 3 used a visual perception task in which participants had to detect brief, peripheral dots while a disk oscillated in the bottom right corner. Full details are reported in Fakche & Dugué (2024, JoCN). **(D)** One period of the oscillation reflects a transition from black to white and back to black again. **(E)** Empirical mode decomposition (EMD) breaks a complex time-series into a finite number of intrinsic mode functions (IMFs). We focus our analyses on the IMFs that arise at the fundamental and second harmonic stimulation frequencies. **(F)** Pipeline for non-Fourier analysis of M/EEG data. The data undergoes standard preprocessing and quality-checking before commencing the main analyses. A peak-locked average is created using peaks in the EEG signal (Exp. 1) or in a photodiode (Exp. 2-3). Solid lines indicate MEG data from one participant. Vertical dotted lines indicate the photodiode peaks. Peak-locking allows us to home in on the rhythmic responses to stimulation without the need for narrowband filtering (which may itself induce rhythmic effects). Peak-locking averaging does not introduce a false sense of rhythmicity (see Supplementary Figure 1). The peak-locked average is decomposed into intrinsic mode functions (IMFs). Grey solid line depicts the IMF at the fundamental frequency; red solid line depicts the IMF at the harmonic frequency; vertical dotted lines indicate the photodiode peaks. Spectral power is computed on the IMFs individually. Spectral power for the harmonic IMF is contrasted to a baseline condition to deduce whether harmonic power is significantly greater than what would be expected without rhythmic sensory stimulation. Grey solid line depicts power for the IMF at the fundamental frequency; red solid line depicts power for the IMF at the harmonic frequency.

We investigate our hypotheses in three datasets that paired two behavioural tasks (see Figure 2A-D) with MEG/EEG recordings and rhythmic light stimulation. Taking the variability in experimental protocols in our stride (see Table 1), we show that rhythmic light stimulation elicits two concurrent but separable neural responses, one at the fundamental and one at the second harmonic, which are spatially and temporally distinct. This is replicable across tasks, electrophysiology recording modality, target frequency band, and is robust to changes in the preprocessing pipeline. Moreover, we demonstrate that the harmonic responses elucidate (1) oscillatory coding of representational content in gamma oscillations, and (2) alpha phase-dependent visual perception. In a second report, we demonstrate how these gamma oscillations impact associative memory retrieval^69^. Altogether, this suggests that multiple oscillatory responses to rhythmic stimulation are a pervasive and robust phenomenon, with each response uniquely impacting perception.

**Table 1.** DiFerences in parameters across experiments. Rather than homogenise the experimental designs, we sought to demonstrate the robustness of subharmonic stimulation across numerous potential mediating variables. This table highlights all variations except preprocessing (see methods for preprocessing pipelines used for each experiment).

| Parameter | Experiment 1 | Experiment 2 | Experiment 3 |
| --- | --- | --- | --- |
| **Final Sample Size** | 31 | 33 | 13 |
| **Task** | Associative memory | Associative memory | Near-threshold perception |
| **Recording Modality** | 128-channel EEG | 306-sensor MEG | 64-channel EEG |
| **Stimulation Frequencies** | 60/40/30/24/20Hz | 68/44/33Hz | 10/8/6/4Hz |
| **Stimulation Waveshape** | Single Frame (~2ms) | Single Frame (~7ms) | Sinusoid |
| **Stimulation Monitor Type** | LCD Monitor | ProPixx Projector | ProPixx Projector |
| **Stimulation Monitor Refresh Rate** | 120Hz | 480Hz | 480Hz |
| **Referencing** | Average of montage | N/A | Mastoids |
| **Control** | No Stimulation | Broadband Flicker | Shuffled Trials |
| **Peak Identification** | EEG Signal | Photodiode | Reconstruction |

## Results

### Rhythmic light stimulation elicits multiple oscillatory responses that are spectrally, spatially, and temporally distinct

We began by asking whether beta-band (20-30Hz) stimulation elicits concurrent oscillatory responses at its fundamental (20-30Hz) and second harmonic (40-60Hz; i.e., gamma) frequencies. Participants completed an associative memory task while undergoing scalp EEG recordings and rhythmic light stimulation. We stimulated at 60Hz, 40Hz, 30Hz, 24Hz, and 20Hz, and included a baseline condition without stimulation. Then, using EMD to unmix overlapping oscillatory signals in the EEG, we focused our analyses on (i) the IMF that most closely matched the stimulation frequency (*f*Hz; from here on, the “fundamental”) and (ii) the IMF which most closely matched the second harmonic (*2f*Hz; the “harmonic”) [see methods for full details on the approach to EMD]. We used the Hilbert-Huang transform to compute the power spectrum for the two IMFs and then compared these power spectra to their equivalents in the baseline condition.

Rhythmic light stimulation significantly increased fundamental power relative to baseline [fundamental; 20Hz: cluster z = 24.96; p < 0.001; 24Hz: cluster z = 16.28; p < 0.001; 30Hz: cluster z = 9.31; p < 0.001; 40Hz: cluster z = 25.80; p < 0.001; 60Hz: cluster z = 12.82; p < 0.001; see Figure 3A-B]. Critically, beta-band stimulation frequencies also led to a significant increase in harmonic power relative to baseline [stimulation at 20Hz, response at 40Hz: cluster z = 1.45; p = 0.043; stimulation at 24Hz, response at 48Hz: cluster z = 7.48; p = 0.002; stimulation at 30Hz, response at 60Hz: cluster z = 8.13; p = 0.001]. These results demonstrate that beta-band rhythmic stimulation produces oscillatory responses at fundamental and harmonic effects that are spectrally separate.

**Figure 3.**
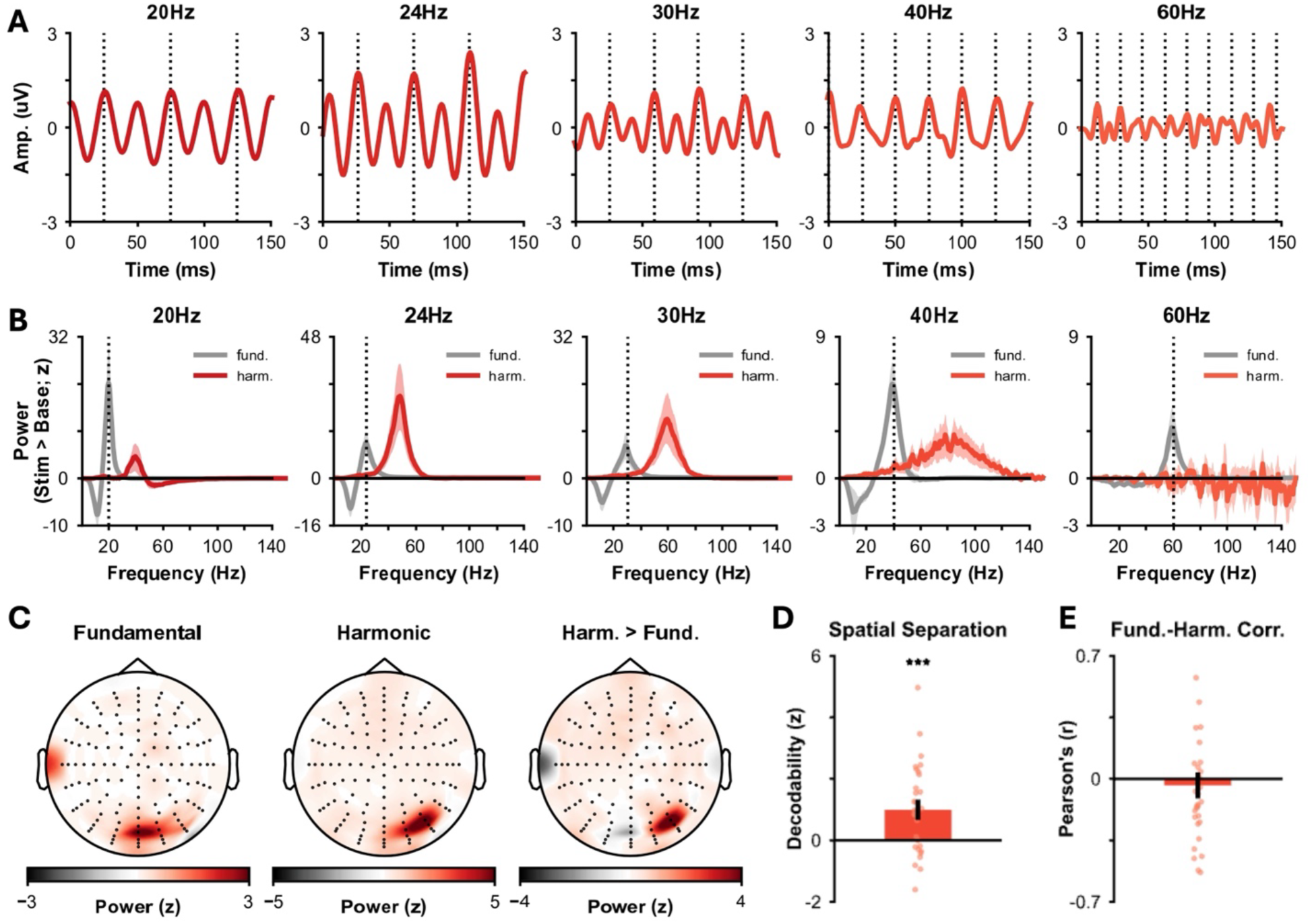
Harmonic EEG responses with the gamma-band following rhythmic sensory stimulation. **(A)** Peak-locked average of data from a single participant for the five stimulation conditions. Clear harmonic responses can be seen following lower-frequency stimulation, but these effects dissipate for higher (>40Hz) stimulation frequencies. Coloured lines depict EEG response averaged across channels. Dotted grey lines indicate approximate time between each stimulation pulse. **(B)** Power for the fundamental and harmonic IMFs, averaged across participants and channels. Coloured lines indicate power for the harmonic IMF; grey lines indicate power for the fundamental IMF; dotted grey line indicates stimulation frequency; shaded areas indicate standard error of the mean. **(C)** Topography of the fundamental (left) and harmonic (middle) IMFs, centred on the fundamental and harmonic frequency respectively and then averaged across participants and conditions. The right topography depicts the diDerence between the two conditions, with positive values indicating greater power for the harmonic IMF and negative values indicating greater power for the fundamental IMF. **(D)** The fundamental and harmonic topographies can be distinguished using linear discriminant analyses (relative to data with shuDled labels). Bar indicates group mean; dots indicate participants. **(E)** No correlation is observed between fundamental power and harmonic power over time. Bar indicates group mean; dots indicate participants.

Notably, harmonics for faster stimulation frequencies (>30Hz) did not show this effect [stimulation at 40Hz, response at 80Hz: cluster z = –0.30; p = 0.557; stimulation at 60Hz, response at 120Hz: cluster z = –1.07; p = 0.878]. Several explanations may account for this, such as the frequency mismatch between the harmonic and endogenous gamma oscillations (∼30-80Hz^21,70,71^) being too great, or that high-frequency oscillations are distorted by the skull^72,73^ (an idea addressed in the MEG data of Experiment 2). Regardless, the lack of harmonic effects in these two conditions rules out the possibility of signal processing decisions (e.g., filtering, re-referencing) artificially introducing the second harmonic during data analysis.

To determine whether these responses emerge from separable neural sources, we used cross-validated linear discriminant analysis (LDA). The classifier was trained to categorise IMF envelopes (fundamental vs. harmonic) using channels as features. We applied the trained classifier to a held-out fold and computed decision values (as in ^74^). We normalised the result against a surrogate distribution generated by repeating this pipeline 20 times with shuffled category labels. This classifier performed significantly better than chance [z = 4.58, p < 0.001; see Figure 3C-D], indicating that the fundamental and harmonic IMFs have distinct topographies.

To determine whether the fundamental and harmonic responses are temporally separate, we correlated the envelopes of the fundamental and harmonic IMFs over time. We found no statistically significant correlation [z = –2.33, p = 0.542s; BF_10_ = 0.229, “moderate” evidence for null^75^; see Figure 3E], suggesting that the temporal dynamics of fundamental and harmonic IMFs are not reliably linked over time. Paired with the topographic analyses above, this suggests that these two neural responses have distinct neural origins.

Altogether, the results from Experiment 1 suggest that beta-band stimulation produces harmonic gamma-band responses that are spectrally, spatially, and temporally distinct from responses at the fundamental frequency.

We conceptually replicated these results in an independent MEG dataset, which used the same task and similar stimulation frequencies (34Hz, 44Hz, & 68Hz; see Figure 4). As above, we found that the slower stimulation frequencies produce harmonic responses within the gamma band [stimulation at 34Hz, response at 68Hz: cluster z = 13.37, p < 0.001; stimulation at 44Hz, response at 88Hz: cluster z = 8.61, p = 0.003] while the fastest stimulation frequency did not [stimulation at 68Hz, response at 136Hz: cluster z = 0.15, p = 0.279]. Aligning with Experiment 1, the 68Hz harmonic response to 34Hz stimulation was spatially and temporally distinct from the 34Hz response (spatial LDA analysis: z = 22.86, p < 0.001; temporal correlation: z = 0.272, p = 0.345; BF_10_ = 0.291, “moderate” evidence for null^75^).

**Figure 4.**
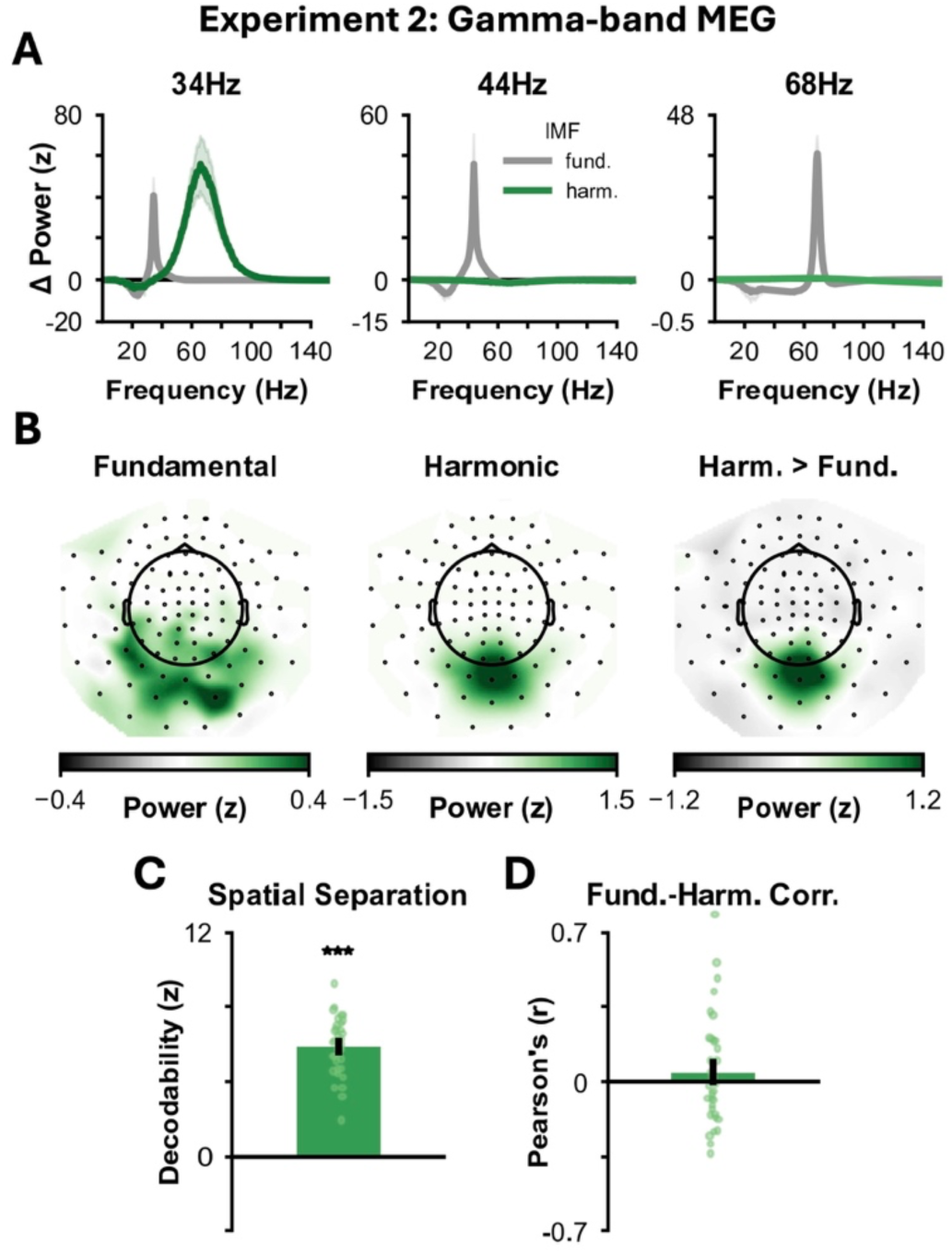
Conceptual replication of harmonic responses in gamma-band MEG and alpha-band EEG data. **(A)** MEG power for the fundamental and harmonic IMFs, averaged across participants and channels. Coloured lines indicate power for the harmonic IMF; grey lines indicate power for the fundamental IMF; dotted grey line indicates stimulation frequency; shaded areas indicate standard error of the mean. **(B)** Topography of the fundamental (left) and harmonic (middle) IMFs, centred on the fundamental and harmonic frequency respectively and then averaged across participants and conditions. The right topography depicts the diDerence between the two conditions, with positive values indicating greater power for the harmonic IMF and negative values indicating greater power for the fundamental IMF. **(C)** The fundamental and harmonic topographies can be distinguished using linear discriminant analyses (relative to data with shuDled labels). Bar indicates group mean; dots indicate participants **(D)** No correlation is observed between fundamental power and harmonic power over time. Bar indicates group mean; dots indicate participants. **(E)** See Panel A. **(F)** See Panel B. **(G)** See Panel C. **(H)** See Panel D. (*** p < 0.001).

The presence of a harmonic effect at 44Hz suggests that some of the higher frequency EEG effects in Experiment 1 may have been attenuated by the skull^72,73^. However, the continued absence of an effect following 68Hz stimulation indicates that the frequency limits of endogenous gamma still mediate the harmonic response, and continues to suggest that harmonics are not attributable to signal processing confounds.

Altogether, these results conceptually replicate those of Experiment 1, demonstrating that fundamental and harmonic gamma-band responses to rhythmic light stimulation are spectrally, spatially, and temporally distinct from beta-band responses at the fundamental frequency.

### Rhythmic stimulation facilitates oscillatory coding in harmonic responding frequencies

If harmonic gamma-band responses reflect changes in endogenous gamma oscillatory activity, these harmonic responses should influence the neural phenomena associated with gamma oscillations. One such phenomenon is oscillatory coding, where incoming visual information is represented in specific phases of gamma oscillations^76–79^. To test this idea, we conducted linear discriminant analysis (LDA) on the MEG data collected while participants were viewing dynamic visual stimuli (see Figure 2A), training the classifier to discriminate the four videos using data from all conditions except 34Hz and applied the classifier to the held-out 34Hz data. We then transformed the resulting decoding time-series into the frequency domain (as in ^80,81^) to determine whether visual information is coded in harmonic gamma-band response.

We first set out to confirm that we could decode visual content in the time-series MEG data. To do so, we compared the time series of LDA decoding to a surrogate distribution built from shuffling video labels, finding that the true time series showed significantly greater decoding than what would be expected by chance (z = 27.85, p < 0.001; see Figure 5A). Then, to pinpoint an oscillatory code, we converted the decoding time-series to the frequency domain and looked for rhythmic fluctuations in representational content. When comparing decoding at the harmonic frequency relative to its neighbours (±5Hz), we observed a significant peak in decoding (z = 2.91, p = 0.006; see Figure 5B-C). We observed no similar effect at 34Hz (z = –1.16, p > 0.5). This suggests that the harmonic gamma-band responses to stimulation rhythmically represent incoming visual information, whereas responses at the fundamental frequency (i.e., beta-band) do not.

**Figure 5.**
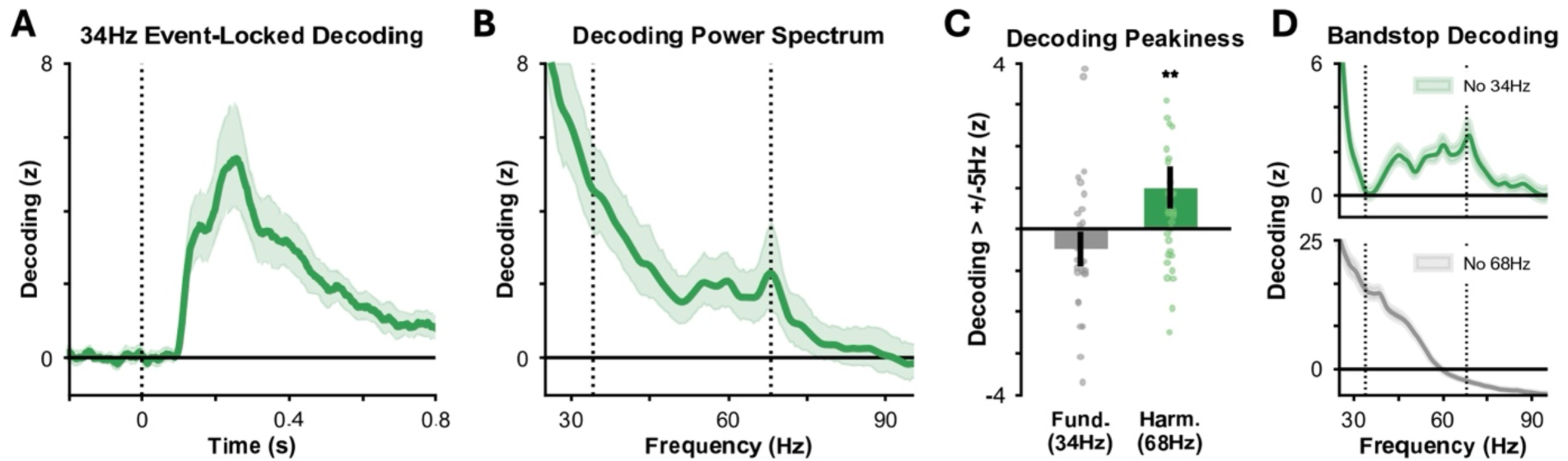
Oscillatory coding of visual content in gamma oscillations. **(A)** Decoding performance locked to stimulus onset (time = 0). Decoding performance reflects the distance to the decision boundary, scaled by the mean and standard deviation of label-permuted surrogate data (1,000 permutations). Solid green line indicates mean performance across participants; shaded area reflects standard error of this mean. **(B)** Decoding performance was transformed to the frequency domain on a trial-by-trial basis and then averaged across participants and trials. Solid green line indicates mean performance across participants; shaded area reflects standard error of this mean. **(C)** Decoding power at 68Hz (the second harmonic) showed significantly greater decoding than neighbouring frequencies at 63/73Hz. No similar effect was observed at 34Hz (the fundamental frequency of stimulation). **(D)** The analyses in Panels B and C were repeated on bandstop-filtered data at 34Hz (top) and 68Hz (bottom). The 68Hz effect persists when using a 34Hz bandstop, but not a 68Hz bandstop.

To rule out the possibility that this effect reflects amplitude differences at 34Hz^82^, we ran two control analyses. First, we bandstop-filtered around 68Hz, reasoning that if 34Hz activity were driving the effect, it should persist when 68Hz activity has been removed. Doing so, however, removed the 68Hz peak in decoding (z = –0.86, p > 0.5; see Figure 5D), suggesting that 34Hz is not driving the effect. Second, we bandstop-filtered around 34Hz to remove potential contributions of 34Hz to the 68Hz condition. Here, the peak at 68Hz remained significant (z = 2.01, p = 0.044; see Figure 5D), further suggesting that 34Hz is not contributing to the effect.

These results demonstrate that gamma-band harmonic responses to rhythmic stimulation support oscillatory coding of incoming visual information in a manner distinct from responses at the fundamental frequency.

### Rhythmic light stimulation elicits multiple oscillatory responses in the theta– and alpha-bands

Confident that beta-band stimulation can reliably induce harmonic responses in the gamma-band that are spatially, temporally, and functionally distinct from responses at the fundamental, we set out to generalise this phenomenon to theta-band stimulation and visual alpha oscillations. We reanalysed data from a recent visual perception study^83^ in which participants underwent EEG while a disk oscillated in luminance on the periphery of participants’ vision (at 4Hz, 6Hz, 8Hz, and 10Hz; see Figure 2C-D). Taking the same approach as in Experiment 1 and 2, we found that rhythmic light stimulation significantly increased fundamental power relative to baseline [4Hz: cluster z = 12.65, p = 0.002; 6Hz: cluster z = 17.63, p < 0.001; 8Hz: cluster z = 14.69, p = 0.002; 10Hz: cluster z = 15.71, p < 0.001; see Figure 6A]. Critically, slower stimulation frequencies produced harmonic responses [stimulation at 4Hz, response at 8Hz: cluster z = 2.51, p = 0.028; stimulation at 6Hz, response at 12Hz: cluster z = 2.97, p = 0.023; stimulation at 8Hz, response at 16Hz: cluster z = 2.81, p = 0.021], whereas faster stimulation at 10Hz did not [stimulation at 10Hz, response at 20Hz: cluster z = –0.296, p = 0.517]. The lack of a 20Hz harmonic response to 10Hz stimulation might seem to contradict past research^55,84^, but we believe this is simply due to a difference in methodology. Previous studies used the FFT to assess harmonic frequencies, meaning the harmonic response could reflect non-sinusoidal properties of the 10Hz oscillation (rather than a true 20Hz oscillation). Our use of EMD sidesteps this issue, and given that we do not see a harmonic response, suggests that harmonic results in these previous studies do indeed reflect non-sinusoidal properties of the 10Hz oscillation rather than a true 20Hz oscillation.

**Figure 6.**
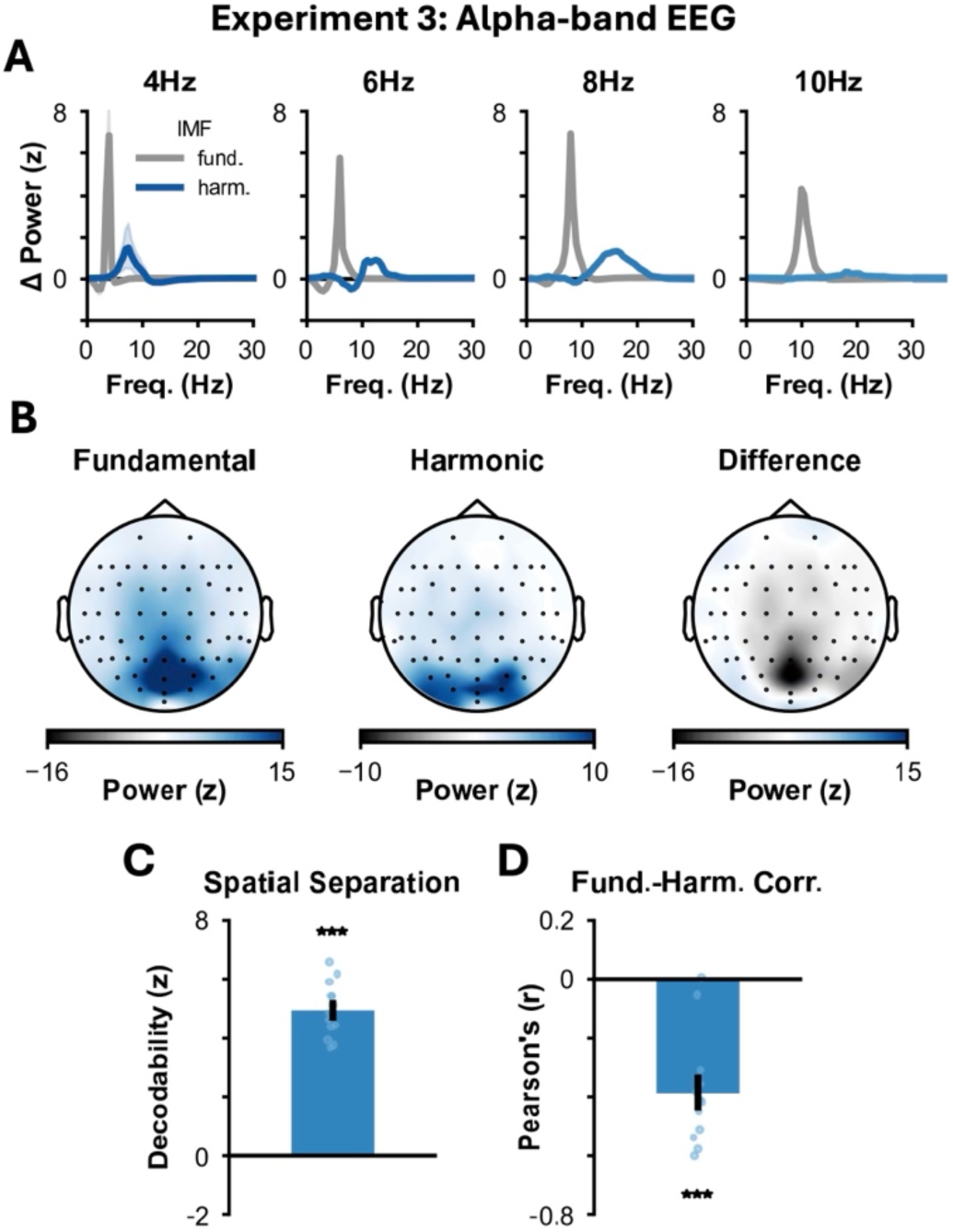
Conceptual replication of harmonic responses in gamma-band MEG and alpha-band EEG data. **(A)** MEG power for the fundamental and harmonic IMFs, averaged across participants and channels. Coloured lines indicate power for the harmonic IMF; grey lines indicate power for the fundamental IMF; dotted grey line indicates stimulation frequency; shaded areas indicate standard error of the mean. **(B)** Topography of the fundamental (left) and harmonic (middle) IMFs, centred on the fundamental and harmonic frequency respectively and then averaged across participants and conditions. The right topography depicts the difference between the two conditions, with positive values indicating greater power for the harmonic IMF and negative values indicating greater power for the fundamental IMF. **(C)** The fundamental and harmonic topographies can be distinguished using linear discriminant analyses (relative to data with shuffled labels). Bar indicates group mean; dots indicate participants **(D)** No correlation is observed between fundamental power and harmonic power over time. Bar indicates group mean; dots indicate participants. **(E)** See Panel A. **(F)** See Panel B. **(G)** See Panel C. **(H)** See Panel D. (*** p < 0.001).

As with Experiments 1 and 2, the fundamental and harmonic IMFs were spatially distinct [z = 18.74, p = 0.001; see Figure 6B-C]. Intriguingly, however, we observed a significant negative correlation between time courses [z = –10.94, p = 0.002; see Figure 6D], suggesting harmonics were strongest when the fundamental response was weakest.

These results, paired with the gamma-band responses above, demonstrate that fundamental and harmonic responses to rhythmic stimulation are replicable and generalisable across frequency bands. Therefore, we suggest that multiplex oscillatory responses to rhythmic input are a pervasive phenomenon in rhythmic stimulation research.

### Individual alpha frequency predicts participant-specific responses at fundamental and harmonic frequencies

Synchronisation theory states that harmonic responses to rhythmic stimulation arise when there is sufficient alignment between the endogenous oscillatory frequency and an integer multiple of the input frequency^40^. Given that endogenous oscillatory frequency differs between individuals, this begs the question: do differences in individual alpha frequency (IAF) explain individual differences in harmonic alpha responses to rhythmic light stimulation?

We began by extracting participant IAFs using eyes-open resting state data (see methods). As no rhythmic stimulation was applied during the resting-state recordings, this means we sidestep the difficulties in extracting IAF that occur when using data involving concurrent stimulation^85^. As the IAFs of the sample were approximately evenly distributed around 9Hz (see Figure 7A), we focused our analyses on the 6Hz stimulation condition to observe a potential interaction across participants where participants with slower IAFs (<9Hz) produce stronger effects at the fundamental (6Hz) and those with faster IAFs (>9Hz) produce stronger effects at the harmonic (12Hz). This interaction would not be observable for other stimulation frequencies, as the IAFs of all participants favoured the second harmonic of the 4Hz condition and the fundamental of the 8Hz and 10Hz conditions (see Figure 7A).

**Figure 7.**
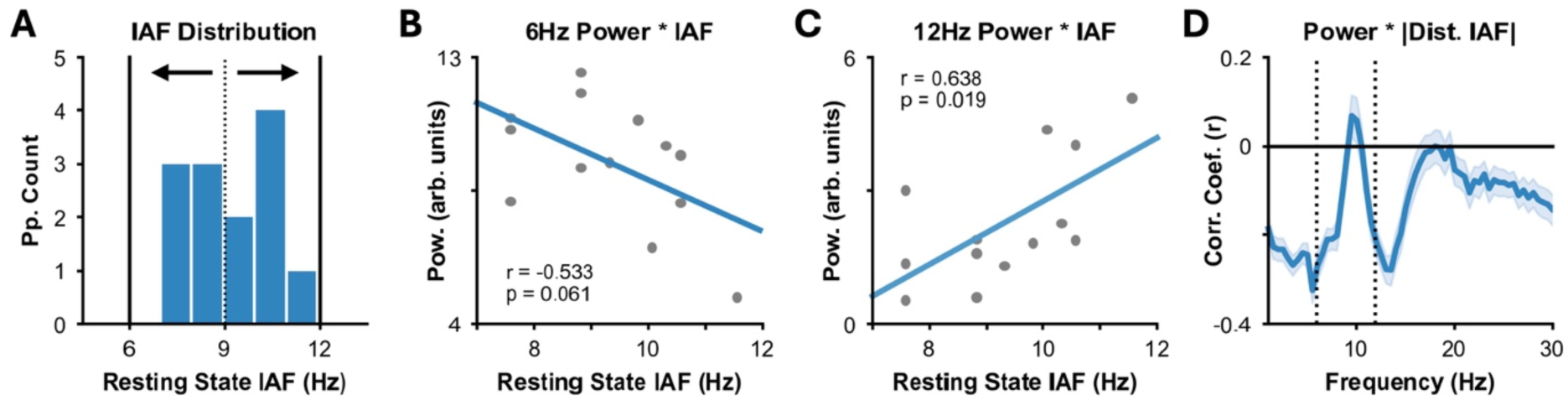
Across-participant variability in harmonic responses relates to individual alpha frequency. **(A)** Individual alpha frequency (IAF) for each participant was calculated using eyes-open resting state data. Following the notion of Arnold Tongue, we hypothesised that 6Hz sensory stimulation would produce a strong response at the fundamental frequency for those with a slower IAF and a greater response at the harmonic (12Hz) for those with a faster IAF. **(B)** We find a negative correlation between IAF and power for the fundamental IMF following 6Hz, indicating that power at the fundamental frequency decreases as IAF increases. Conversely, we find a positive correlation between IAF and power at the harmonic IMF, indicating that harmonic power increases as IAF increases. Dots indicate individual participants. Lines reflect line-of-best-fit computed using least-squares regression. **(C)** Magnitude of correlation between power and the absolute distance to the IAF across all frequencies. At both 6Hz and 12Hz, there is a significant negative correlation indicating that participants with IAFs further away from 6/12Hz show reduced power at these frequencies. **(D)** There is no effect at intermediate alpha frequencies (i.e., 7-11Hz) indicating that this is not a trivial correlation between IAF and alpha power but an effect specific to the fundamental and harmonic frequencies of 6Hz stimulation. Dark blue line indicates mean correlation coefficient across channels; shaded area indicates standard error of correlation coefficient across channels.

When participants underwent 6Hz stimulation, we observed a significant positive correlation between IAF and 12Hz power (r = 0.638, p = 0.019), indicating that participants with faster IAFs produced stronger harmonic responses. Conversely, when correlating IAF and 6Hz power, we observed a trending negative correlation (r = –0.533, p = 0.061; see Figure 7B-C), indicating that participants with slower IAFs produced stronger responses at the fundamental. No similar effects were observed for power at frequencies other than the fundamental or harmonic (see Figure 7D). These results indicate that harmonic responses to rhythmic light stimulation are strongest when the second harmonic of the rhythmic input matches the fundamental of the endogenous oscillatory frequency.

The absence of resting state data for the other datasets precluded similar analyses for individual gamma frequencies. However, descriptively speaking, we can observe a related phenomenon in Experiment 1 when stimulating at lower frequencies (i.e., 20Hz; see Supplementary Figure 3). Given that the second harmonic (40Hz) sits at the lower bounds of the canonical visual gamma band (∼30-80Hz^21,70,71^), we might expect participants with fast endogenous gamma to show responses at higher harmonics (e.g., 3*f*, 60Hz). Visual inspection of the peak-locked averages supports this, with several participants showing clear harmonics at 60Hz, rather than 40Hz (for examples, see Supplementary Figure 3A). Moreover, we observed a negative correlation between harmonic power at 40Hz and 60Hz across participants (r = –0.677, p < 0.001; see Supplementary Figure 2B), suggesting participants show responses at either 40Hz or 60Hz but not both. These results suggest that harmonic responses to 20Hz stimulation are influenced by the resonant frequency of endogenous gamma oscillations.

It is worth noting that the analyses in this subsection were between-participant and conducted on a relatively small sample (Experiment 3: n=13), which likely overestimates the effect size. Therefore, it will be of interest to see how these results scale in larger datasets. Nonetheless, these observations suggest that harmonic responses to rhythmic stimulation depend upon the participant-specific resonant frequency of endogenous alpha/gamma oscillations.

### Co-occurring harmonic and fundamental neural responses separably influence human visual perception

Finally, we asked whether harmonic responses to rhythmic stimulation impact human behaviour by investigating whether the phase of harmonic alpha-band oscillations impacts visual perception^19,20,83,86,87^.

To address this, we expanded upon the original analysis of Experiment 3, which demonstrated that visual perception fluctuates according to the phase of the fundamental frequency of stimulation^83^. We binned behavioural data according to the phase of rhythmic light stimulation and then used sine waves to model this behavioural time-series. We built two models: a “simple” model consisting of a single sine wave at the fundamental frequency and a “complex” model consisting of two sine waves, one at the fundamental frequency and the other at the second harmonic (see Figure 8A-B).

**Figure 8.**
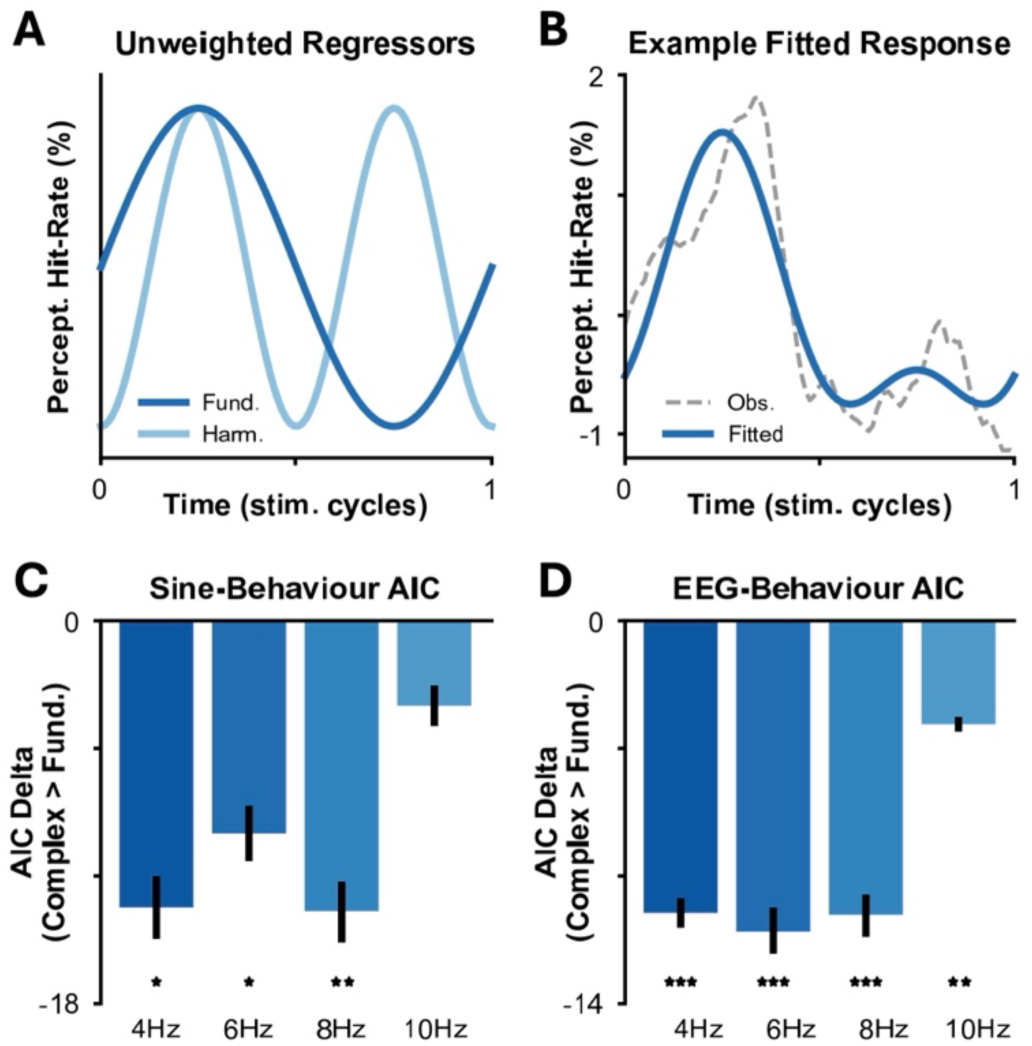
Fundamental and harmonic neural responses interact to impact visual perception. **(A)** We modelled perceptual hit rate (i.e., the behavioural responses) over time as the weighted sum of two sine waves – one at the fundamental frequency (dark blue) and one at the second harmonic (light blue). **(B)** Example fit to behavioural data. Dotted grey line indicates perceptual hit rate over the course of a stimulation cycle. Solid blue line indicates the fitted sine waves. **(C)** To identify whether how the two sine waves impacted modelling fitting, we compared the Akaike Information Criterion (AIC; left) and Bayesian Information Criterion (BIC; right) for a simple model using only the sine wave at the fundamental frequency against a complex model using both sine waves. Aside from 10Hz, we find a significant improvement in model fit when using the complex model. Bars indicate mean AIC/BIC across participants, error bars indicate standard error in AIC/BIC across participants. **(D)** Example model fit using IMF waveshapes in place of sine waves. When fitting the models using IMF waveshapes in place of sine waves, we replicate our results, indicating that neural responses to stimulation at fundamental and harmonic frequencies interact to modulate perceptual performance. ** p < 0.01; * p < 0.05.

We used Akaike Information Criterion (AIC) to compare the two models; if AIC decreases for the complex model, it means that the harmonic sine wave improves the fit to behavioural data relative to what the fundamental sine wave achieved in isolation (after accounting for the difference in the number of free parameters). Indeed, we found that the “complex” model explained behavioural performance to a significantly greater degree than the “simple” model for all stimulation conditions except 10Hz (i.e., the one condition that did not produce a harmonic response; 4Hz: z = –5.24, p = 0.019; 6Hz: z = –5.15, p = 0.006; 8Hz: z = 5.78, p = 0.004; 10Hz: z = –9.21, p = 0.063; see Figure 8C). These results suggest that the phase of harmonic responses impacts visual perceptual performance in a way that is complementary, but separable, to responses at the fundamental frequency.

To tie this back to the putative alpha oscillations in the EEG, we refined the above analysis by replacing the sine waves with participant-specific waveshapes (extracted from the fundamental and harmonic IMFs). Again, we found that the “complex” model explained behavioural performance to a significantly greater degree than the “simple” model (AIC; 4Hz: z = –21.24, p < 0.001; 6Hz: z = –17.22, p < 0.001; 8Hz: z = –19.62, p < 0.001; 10Hz: z = –4.65, p = 0.004; see Figure 8D).

To rule out the possibility that AIC did not sufficiently penalise the “complex” model, we ran a control analysis using phase-shifted versions of the harmonic IMF, finding that the true “complex” model better explained behavioural changes in performance than any other phase-shifted “complex” model (see Supplementary Figure 4).

These results indicate that harmonic alpha activity induced by theta-band stimulation can impact near-threshold visual perception. This harmonic effect is summed with the effect from neural responses at the fundamental frequency, indicating that fundamental theta-band and harmonic alpha-band responses to stimulation have separable causal effects on behavioural performance.

## Discussion

We demonstrate that rhythmic light stimulation elicits multiple concurrent neural responses that have separable impacts on human perception. Using EMD^66,67^, an approach that distinguishes multiplexed oscillations from non-sinusoidal waveforms^26,57–59^, we show that rhythmic stimulation elicits harmonic responses within the canonical gamma and alpha bands in the visual cortex. These responses influence human perception in a manner distinct from co-occurring responses at the fundamental frequency, with gamma-band responses contributing to oscillatory coding^21,76,77^ and harmonic alpha responses contributing to phase-dependent visual perception^19,20,83,86,87^ (see also ^69^ for how harmonic gamma-band responses impact associative memory). We observe these effects across three independent datasets, two distinct tasks, two recording modalities, and multiple stimulation frequencies, using a variety of preprocessing pipelines, stimulation parameters, stimulation hardware (i.e., monitors/projectors), and baseline conditions. This steps beyond past work that has demonstrated the presence of FFT-derived harmonics in electrophysiology^60–65^ by revealing the nature of these harmonics: namely, that these reflect multiple, multiplexed oscillatory responses, rather than simply reflecting harmonic components of a single, non-sinusoidal oscillation. We propose that harmonic oscillatory responses to rhythmic light stimulation are a reliable and pervasive phenomenon that influences human perception, cognition, and behaviour in ways unique to neural responses at the fundamental frequency. More generally, we suggest that oscillatory responses to rhythmic brain stimulation (and the associated behavioural outcomes) are more complex than is often acknowledged.

### “Neural side eCects” in neuromodulation

Most rhythmic stimulation experiments seek to match stimulation frequency to a specific oscillatory target. We demonstrate that this approach may unintentionally modulate other oscillations that have a harmonic relationship with the stimulation frequency. Therefore, these harmonic effects (and their behavioural consequences) could be considered “neural side effects”. This provides a possible explanation of variability in rhythmic stimulation studies: unaccounted neural responses introduce a series of frequency– and participant-specific confounds that bias results and interpretations. Accounting for these “side effects”, therefore, may enhance the accuracy and precision of studies investigating how rhythmic stimulation influences human perception, cognition, and behaviour, leading to more conclusive evidence for causal mechanisms and improved efficacy of neural interventions.

We envisage that these effects could be accounted for analytically and/or methodologically. First, it can be handled analytically, as we have done above: decomposing the recorded MEG/EEG signal into intrinsic mode functions (IMFs) and incorporating them into a single model that aims to explain behaviour (see Figure 8). This modelling approach means the IMFs cannot explain the same variance in behaviour, and therefore helps dissociate their contributions to changes in behaviour. On a therapeutic level, however, it may be better to tackle this issue methodologically, minimising harmonic neural responses to stimulation by modifying the stimulation protocol itself. Empirical demonstrations of such methods are uncommon, but modelling work suggests that adding white noise to the stimulation period may be able to suppress neural responses beyond the fundamental frequency^88^. Notably, however, suppressing the harmonic response may undermine an intervention if the harmonic response is the sole driver of the observed cognitive change (e.g., gamma oscillatory coding; Figure 5). Therefore, one needs to take careful consideration of the strategy used to account for the harmonics so that they can maximise interpretability and impact without abolishing the effect of interest.

### Entrainment, resonance, or steady-state visually-evoked responses?

We show that fundamental and harmonic responses to rhythmic stimulation differ in spectral, spatial, and temporal dynamics. But do they also differ in how they are generated?

There is an ongoing debate about the neural mechanisms that generate electrophysiological responses to rhythmic stimulation^6,47,89,90^. An electrophysiological signature of rhythmic stimulation can arise when (i) an endogenous oscillator adjusts its frequency and phase to match the input (*entrainment*^2^); (ii) an endogenous oscillating response resets with each stimulation pulse (*resonance*), or (iii) aperiodic steady-state visually-evoked potentials (*SSVEPs*) are superimposed^47^. Distinguishing between these mechanisms is crucial for both fundamental and translational research as the underlying mechanism dictates how rhythmic stimulation impacts cognition. For example, Alzheimer’s disease is associated with disrupted gamma oscillations^25^, but if gamma-band stimulation only elicits SSVEPs (rather than modulating dysfunctional gamma oscillations), the intervention would be unlikely to tackle the underlying cause of the problem.

While it remains ambiguous whether a rhythmic response at the fundamental frequency can reflect entrainment^6,47,89,90^, we suggest that the harmonic responses observed here inherently reflect entrainment because:

1) The harmonic response frequency is always an integer multiple of the fundamental frequency (that is, the frequency of the harmonic response shifts with the input frequency, as in entrainment). Resonance, in contrast, would predict that the higher frequency response would have a fixed numerical value matching the resonant frequency of the endogenous rhythm.
2) There are two cycles of the harmonic response per stimulation pulse, making it difficult to attribute this to SSVEPs. One could argue that harmonics reflect a reversal^3,91^, with SSVEPs occurring at both pulse onset and offset, but Experiments 1 and 2 use asymmetric stimulation protocols (Figure 2B), so reversals cannot explain the equidistant peaks observed in peak-locked averages of these experiments (Figure 3A and Supplementary Figure 3A). Moreover, reversals cannot explain why some participants showed responses at the third harmonic (Supplementary Figure 3A).
3) The harmonic responses are strongest when their frequencies align with endogenous resonant frequencies, and vary across participants based on individual differences in resting oscillatory frequency. This dependence on endogenous oscillatory alignment is a characteristic of entrainment, but not of SSVEPs nor resonance.

Therefore, we hypothesise that harmonic responses reflect entrained oscillations. A confirmatory test of this hypothesis could involve investigating the persistence of oscillatory responses after the offset of stimulation^39,92,93^ (i.e., “oscillatory echoes”): if the oscillatory responses persist at the exact harmonic frequency, it would provide strong evidence for entrainment. If, in contrast, oscillatory responses either immediately shift to a neighbouring harmonic frequency or disappear entirely, responses might better be ascribed to resonance or steady-state evoked responses. Unfortunately, in all our experiments, the offset of the stimulation coincided with the onset/change of other facets of the task (e.g., the onset of a response prompt), introducing a source of covariance that precludes the testing of the hypothesis here.

The responses at the fundamental which we observe here present the same interpretative challenges as previous work^6,47,89^. Since fundamental responses appear across all stimulation frequencies, SSVEPs likely contribute significantly to their generation. This hypothesis is supported by the observation that fundamental responses diminish in power when harmonically responding alpha activity increases, mirroring reductions in early visually-evoked responses during periods of high alpha activity^94^. However, this does not rule out the possibility of entrainment also contributing to the fundamental response (e.g., ^38^).

In sum, harmonic responses most parsimoniously align with the concept of entrainment, whereas fundamental responses remain ambiguous. As it appears easier to relate harmonic responses to entrainment, we propose that future entrainment research should focus on harmonic responses to circumvent pervasive questions about the mechanisms underlying rhythmic electrophysiological responses, which in turn will enable us to arrive at stronger and clearer conclusions about how neural oscillations causally impact perception, cognition, and behaviour.

### Causal role of alpha phase in visual perception

There is considerable interest in the role of alpha oscillations in visual perception. Many correlative studies suggest that perceptual behaviour depends on the phase of alpha^19,20,83,86,87^, but results are not always consistent^95,96^ with evidence indicating that these effects may be mediated by other phenomena^97^ (e.g., oscillatory power^20,98^; cross-region interactions^87^). Causal studies have aimed to establish the primacy of visual alpha oscillations by directly manipulating these rhythms to modulate behaviour^83,99–104^. While such approaches do modulate behaviour, these studies are susceptible to confounds such as phase-dependent phosphenes or luminance changes that impact vision. This leaves the possibility that stimulation-induced changes in neural activity reflect SSVEPs rather than modulations of true oscillations. Harmonic responses to stimulation, however, circumvent these concerns and, therefore, may be key to resolving this debate.

We show that fundamental and harmonic responses to theta-band stimulation have complementary influences on visual perceptual behaviour. As the fundamental and harmonic phases are orthogonal, luminance differences at the fundamental frequency cannot explain the harmonic effect. Moreover, as discussed above, the harmonic response cannot be attributed to an SSVEP. Therefore, the contributions of the harmonic effect are best ascribed to modulations of endogenous alpha oscillations, supporting the case for a causal role of alpha oscillatory phase in visual perception.

### Gamma oscillations and oscillatory coding

Incoming visual sensory information is thought to be represented in a gamma oscillatory code^76–79,105,106^. We find support for this idea in Experiment 2, where we observed that representational fidelity of stimulus-specific visual information rhythmically fluctuated in accordance with the gamma-band harmonic responses to 34Hz stimulation. Notably, the fundamental response showed no effect, indicating a dissociation between fundamental and harmonic responses in the representation of sensory information. These results further support the theory that harmonic responses reflect changes in true oscillatory activity which contribute to neural processing distinct from responses at the fundamental.

The detection of gamma-band oscillatory codes is challenging to test in humans as (i) gamma frequencies overlap with muscle artifacts^72,73^, introducing noise into measures, (ii) there is substantial variability in gamma frequencies across participants^49,90,107^, complicating group-level analyses, and (iii) stimulus properties impact the exact frequency of the gamma response^108^, introducing across-trial variability. We show that rhythmic light stimulation can circumvent these issues by amplifying the code at a known frequency, both across participants and across naturalistic visual stimuli. This offers a new means to test neuroscientific hypotheses about oscillatory coding that may otherwise have proven impossible to assess.

### Alternative explanations

Harmonic responses to stimulation are often viewed as artifacts^3,26,26,58,59^. However, we believe that oft-cited reasons for doing so, reviewed in turn below, have no bearing on our findings:

- Fourier transform artifacts: Applying a Fourier transform to non-sinusoidal oscillations produces harmonics at multiple frequencies^26,57–59^, so harmonic effects observed using this approach are often assumed to be artifacts. We circumvent this issue by using EMD^66,67^, which disentangles multiplex harmonic responses from non-sinusoidal fundamental responses^68^. Critically, we show that harmonic responses to stimulation act in differing ways from fundamental responses on both a neural and behavioural level – a result that would not occur if the harmonic response were a reflection of a non-sinusoidal response at the fundamental. Further evidence to suggest that our effects do not reflect non-sinusoidal artifacts include (i) the observation of a singular harmonic response (typically at the second harmonic, but see Supplementary Figure 3) which differs from non-sinusoidal artifacts (which produce responses at multiple harmonic frequencies^3^) and (ii) the observation that harmonic effects only arise when they approximate with endogenous oscillatory frequencies, which differ from non-sinusoidal artifacts which produce harmonics regardless of stimulation frequency^55^.
- Stimulation reversals: Reversals occur when the brain elicits evoked responses to both the onset and offset of the stimulation pulse. If the “on” and “off” durations match, second harmonics can be attributed to reversals (e.g., ^3,91^). However, this was not the case in Experiments 1 and 2, where the pulse was only presented for a single frame (8.33ms in Experiment 1; 2.08ms in Experiment 2; see Figure 2B), meaning reversals are unlikely to explain these effects. Moreover, reversals fail to account for participants showing harmonic responses at the third harmonic (i.e., three oscillatory cycles per pulse; see Figure 5E) or why harmonic responses only arise when they align with endogenous oscillations. More generally, the high frequency of these reversals (i.e., 120Hz in Experiment 1; 480Hz in Experiment 2) means they will likely be decimated by the low-pass filtering properties of the thalamus^39,109^. This means any reversal effect, or more general “on-off” effect relating to non-linear stimulation waveshapes^3^, would not reach the occipital sources of our main effects.
- Filtering artifacts: Narrowband filtering can distort stimulation-evoked responses to create spurious oscillations^110^. We mitigated this by using peak-locked averages in place of narrowband filters. Furthermore, filtering artifacts cannot explain why, when using the same filters for all data, we see inter-subject and frequency-dependent variability in harmonic responses to stimulation.
- Dipoles: Second harmonics could be explained by the EEG picking up mirrored activity from dipoles (that is, the fundamental and harmonic responses are two sides of the same coin). However, this cannot explain (i) why the fundamental and harmonic responses have distinct effects on visual perception and oscillatory coding; (ii) why the fundamental and harmonic responses are uncorrelated over time in Experiment 1 and Experiment 2; and (iii) why some participants show harmonic responses at the third harmonic (see Supplementary Figure 3).

Therefore, our results indicate that the effects reported here are genuine harmonic responses to stimulation, distinct from those at the fundamental response, which should not be indiscriminately dismissed as artifacts.

### Future directions

Our findings question the idea of a one-to-one mapping between rhythmic stimulation and neural response. Instead, they suggest that rhythmic input elicits a multitude of neural responses, but much remains to be explored regarding how these multiplex responses influence cognition, including:

- *Data-driven detection of neural responses:* We adopted a hypothesis-driven approach to defining the neural responses within our transfer functions, but what if this is not feasible? In such instances, data-driven approaches such as generalised eigendecomposition (GED) or bicoherence may offer solutions. For example, GED can isolate a principal neural response to rhythmic stimulation^111^ and separate harmonic components of an FFT^112,113^. Given the use of FFT in these instances, it remains unclear whether these harmonic components reflect distinct neural oscillations or the non-sinusodiality of a single oscillation. However, integrating GED with approaches such as EMD may enable data-driven ways to capture multiple, separable oscillatory responses to rhythmic stimulation.
- *Causal chains:* Our analysis assumed that the fundamental and harmonic responses to rhythmic stimulation occur independently. However, it remains possible that one neural response impacts another. Such interactions are hinted at by the negative correlation we observe between harmonic alpha power and fundamental theta responses, and by other studies which stimulate one region with the aim of having knock-on effects in task-critical regions^114–116^. Accounting for these interactions would further clarify the causal roles of different neural responses in cognition.
- *Subharmonics:* We focused on harmonic neural responses to rhythmic stimulation as this is well-documented in physics^40^ and biology^42,43,45,117^, but could this principle generalise to subharmonics? Previous studies cast a light on this, showing that 130Hz stimulation of the basal ganglia modulates 65Hz gamma activity^118–120^. Open questions remain about whether this utilises the same mechanisms for generation as harmonic responses. Nonetheless, these results add further support to our claim that a single rhythmic stimulation protocol can elicit a complex array of neural responses.
- *Generalisability to TMS/tACS:* We postulate that transcranial magnetic stimulation (TMS) and transcranial alternating current stimulation (tACS) will produce similar effects to what we observe with rhythmic light stimulation. Indeed, theta-band tACS elicits alpha-band harmonics in ferrets^121^. Moreover, as responses to rhythmic light stimulation are likely weaker than those elicited by TMS/tACS, and as harmonic responses to rhythmic inputs require a strong input^40^, one might hypothesise that harmonic responses may be more pronounced in TMS/tACS. In other words, our effects may be understating rather than overstating the influence of harmonic responses on neural and behavioural outcomes.
- *Precision and personalised neurostimulation:* Accounting for complex neural responses in future research and clinical designs can improve the reliability of scientific findings and efficacy of clinical treatment. For example, developing stimulation protocols that maximise the target neural response while minimising other neural “side-effects” (e.g., ^88^), or including additional control conditions (e.g., harmonic frequencies) could provide greater insight into the mechanisms underpinning neural and behavioural changes.

### Summary

Rhythmic light stimulation (and neuromodulation more broadly) plays a critical role in translating fundamental neuroscience into clinical applications. However, the results of such studies are often highly variable, limiting the development of effective treatments for cognitive and neurological disorders. Here, we show that neural responses to rhythmic light stimulation are more complex than is often acknowledged. Specifically, rhythmic light stimulation elicits multiple neural responses with distinct spectral, spatial, and temporal profiles, each exerting separable influences over human behaviour and varying across individuals. Given that complex neural responses to neuromodulation appear to be the rule, not the exception, we propose it is critical to account for the myriad neural responses that translate rhythmic inputs into behavioural outcomes if we are to advance fundamental and clinical neuroscience. Indeed, a deeper understanding of this phenomenon may well reshape how we approach neuromodulation in the lab, in the clinic, and beyond.

## Methods

### Experiment 1

#### Participants

Thirty-two participants were recruited (mean age = 20.1 years; age range = 18-26 years; 71.9% female; all right-handed [self-reported]). Participants were compensated with course credit or financial reimbursement. Participants provided informed consent before starting the experiment. Ethical approval was granted by the Research Ethics Committee at the University of Birmingham.

#### Behavioural task

Participants completed a paired-associates episodic memory task, learning 192 video-word pairs across 3 blocks. During encoding, each trial began with a fixation cross (1 ± 0.2 s jitter), followed by a video clip and then an English noun (both presented for three seconds). There were four videos in total, each paired with 48 nouns. Videos were presented in a pseudo-randomised order such that each video would appear twice in a series of eight trials. After encoding the video-word pair, participants reported whether they perceived the screen to be flickering during stimulus presentation. Participants had three seconds to respond using the keyboard before the next trial began.

After being exposed to all pairs of a block, participants were then subjected to a retrieval test. During retrieval, each trial began with a fixation cross (1 ± 0.2 s jitter), followed by an English noun (presented for three seconds). These nouns were taken from the immediately preceding encoding phase. Participants had three seconds to select which of the four videos they thought was associated with the word. Following this, a prompt asked whether the participant felt they were confident in their response. Again, participants had three seconds to provide a binary answer (“yes” or “no”).

#### Rhythmic light stimulation protocol

Rhythmic light stimulation was delivered using a 120Hz LCD monitor. Participants sat approximately 50cm away from the monitor. For this, a black box was superimposed over target stimuli and fixation crosses during both encoding and retrieval. The box would appear for a single frame (8.33ms; see Figure 2B) of every cycle of the target frequencies (15Hz, 17Hz, 20Hz, 24Hz, 30Hz, 40Hz, 60Hz). The highest stimulation frequency (60 Hz) surpassed the flicker fusion threshold and appeared as a static grey box. The control condition, where no flickering occurred, used a static grey box to perceptually match the 60Hz condition. Due to difficulties separating the 15Hz and 17Hz conditions for endogenous alpha activity, these trials were not included in the main analyses.

There were a total of 24 trials per stimulation before preprocessing. This was sufficient to determine neural differences between stimulation conditions and the control condition, but lacked power for linking to cognition and behaviour. This was remedied in Experiment 2, which reduced the number of stimulation conditions to boost trial count per condition.

#### EEG acquisition and preprocessing

EEG was recorded from 128 Ag/AgCl active electrodes using a BioSemi Active-Two amplifier (sampling rate = 1,024 Hz). Data were preprocessed using MNE Python^122^ following the FLUX pipeline^123^ with minor adaptations for EEG. The EEG was bandpass filtered (1–256 Hz) and notch filtered (50 Hz, 100 Hz, 150 Hz, 200 Hz, and 250 Hz). Muscle artifacts were detected automatically using MNE’s *annotate_muscle_zscore*, identifying activity between 70–125 Hz exceeding a z-score of 10. Bad channels were identified using RANSAC^124^. Following this, independent component analysis (ICA) was applied to remove spatially stable artifacts (e.g., blinks, saccades). Muscle artifact and bad channel detection were repeated after ICA to remove residual noise. Bad channels were then interpolated using the average of neighbouring electrodes.

EEG data were epoched around stimulus onset (encoding: video onset; retrieval: noun onset), starting 2 seconds before stimulus onset and ending 2 seconds after stimulus offset.

#### Participant exclusion

To maximise signal-to-noise, participants who failed to show the prototypical rhythmic responses at the fundamental frequency were excluded from the main analyses. To identify these participants, for each participant and trial, Morlet wavelets were applied to the preprocessed data (width = wavelet frequency / 2). SpecParam^125^ was used to remove the aperiodic slope from the resulting power spectrum. To isolate the peak at the fundamental frequency, power at the fundamental frequency was normalised against power at +/-1Hz the fundamental frequency (e.g., for the 24Hz condition, power at 23Hz and 25Hz was averaged and then subtracted from power at 24Hz). A cluster-based permutation test (permutations = 1,000; conducted across trials, for each participant individually) was used to determine whether a participant showed a reliable peak at the fundamental frequency (i.e., when p < 0.05). One participant did not pass this threshold, leaving thirty-one participants for further analysis.

#### Spectral separation analyses

To visualise the complex waveforms elicited by rhythmic stimulation, EEG data were first peak-locked (Figure 3A). The data were high-pass filtered at 15 Hz to attenuate low-frequency activity (e.g., endogenous alpha oscillations) and z-scored across time and channels to normalise scales across participants. Peaks in the EEG signal exceeding a prominence threshold of 0.5 standard deviations were detected and used as markers for re-epoching: each new epoch began 0.1 second after the detected peak and lasted 1 second. By excluding the first 0.1 s after the peak, we minimised aperiodic contributions to the peak-locked average. Peak-locked epochs were split by stimulation condition and averaged separately for every condition-channel pair.

Empirical mode decomposition (EMD) was then applied to the peak-locked averages to extract intrinsic mode functions (IMFs). For each channel, an iterated masked sift^68^ was used to extract five IMFs. We elected to extract five IMFs as we found no additional benefit to including further IMFs, and the inclusion of each IMF ramps up computational time. While five IMFs might seem low for EEG-based analyses, it is worth acknowledging that EMD was applied to the peak-locked average, not the raw time series. This averaging procedure leads to a small number of clearer rhythmic components (see Figure 3A; Supp. Fig. 3A), making searching for more than five IMFs redundant. Indeed, increasing the number of IMFs extracted from the peak-locked averages had no impact on the significance of the results. The instantaneous frequency, amplitude, and phase of each IMF were computed using the normalised Hilbert transform. The Hilbert-Huang transform was then used to derive the power spectrum from the instantaneous frequency and amplitude data. The fundamental IMF was defined as the IMF whose mean instantaneous frequency was closest to the stimulation frequency. The harmonic IMF was the IMF whose mean instantaneous frequency was closest to twice the stimulation frequency.

For each participant, the resulting power spectra were standardised using the standard deviation across channels and conditions. Cluster-based permutation tests (1,000 permutations) assessed whether power at the fundamental and harmonic frequencies following stimulation differed from the baseline (i.e., no stimulation) condition. Clusters were considered significant when p < 0.05.

#### Spatial separation analyses

Here, we restricted analyses to the conditions that showed significant effects at both the fundamental and harmonic frequencies (i.e., 24Hz, 30Hz) as there was little sense in searching for spatial differences in conditions which found no evidence for multiple neural responses. Peak-locked averaging and IMF power were computed as before, but done separately for each trial rather than averaging across trials within a condition. We pooled trials across stimulation conditions to produce two categories: one for fundamental power and one for harmonic power.

To examine topographic differences between fundamental and harmonic responses, a five-fold cross-validated linear discriminant analysis (LDA) was used. Four folds formed the training set, with the fifth serving as the test set. Decision values were computed for each trial of the test set^126^, which were then averaged across trials. This process was repeated 1,000 times using shuffled labels to generate a surrogate distribution of chance-level decision values. The true decision value was then z-scored using the mean and standard deviation of the surrogate distribution.

A one-sample permutation test (1,000 permutations) was conducted to determine whether z-scored decision values, pooled across participants, significantly differed from the chance-level surrogate distribution (p < 0.05).

#### Correlations between fundamental and harmonic responses

To test whether fundamental and harmonic responses co-varied, fundamental and harmonic responses (as computed for the spatial separation analyses) were averaged across channels. A Pearson correlation was conducted across trials for each participant separately. The resulting correlation coefficients were then pooled across participants and subjected to a one-sample permutation test (as in the spatial separation analyses; significance: p < 0.05). To aid interpretation of null effects, one-sample Bayes Factor t-tests were used (as implemented in JASP^127^).

### Experiment 2

This experiment largely matches Experiment 1. For brevity, we only describe the methods and details that differ between Experiment 1 and 2.

### Participants

37 participants were recruited (mean age = 24.7; age range = 18-37; 64.9% female; all right-handed [self-reported]).

### Experimental task

See Experiment 1.

### Rhythmic light stimulation protocol

Light stimulation was delivered using a 1,440Hz ProPixx projector at stimulation frequencies of 34Hz, 44Hz, and 68Hz. Participants sat approximately 120cm away from the monitor. In addition to a baseline condition that matched Experiment 1, we included a second baseline that used a broadband 14-54Hz flicker (centred on 34Hz).

### MEG acquisition and preprocessing

MEG was recorded using a 306-channel MEGIN Elekta Triux system, with a 1,000Hz sampling rate. The raw data were corrected using Maxwell filters, with bad channels being marked and removed. Data was then bandpass filtered between 0.5Hz and 220Hz, with notch filters at 50Hz, 100Hz, 150Hz, and 200Hz to attenuate line noise. Muscle artifacts were detected automatically using MNE’s *annotate_muscle_zscore*, identifying activity between 110Hz and 140Hz that exceeds a z-score of 10. ICA and epoching were conducted as in Experiment 1.

### Participant exclusion

Four participants did not pass the inclusion threshold (as described in the methods for Experiment 1), leaving thirty-three participants for further analysis.

### Spectral separation analyses

Analyses matched that of Experiment 1, with three exceptions.

1. Rather than detect peaks in the MEG data itself, we used data from a photodiode that was recorded simultaneously with the MEG. The photodiode provides the flicker ground truth, so it is theoretically the better choice for peak detection.
2. Magnetometers and gradiometers were combined between computing the power spectra and conducting statistical analyses. As the magnetometers and gradiometers were already placed in a common unit space through normalisation, their combination simply reflected taking the mean between the three sensors (one magnetometer, two gradiometers) recording from the same position.
3. We contrasted the periodic stimulation conditions against the broadband stimulation baseline rather than a stimulation-free baseline. This provided us with an “active” control that rules out the possibility that light stimulation produces harmonic oscillatory regardless of whether the stimulation protocol itself is rhythmic or aperiodic.

### Spatial separation analyses

Analyses matched Experiment 1, with the exception that the trials used came from the two significant conditions: 34Hz and 44Hz.

### Correlations between fundamental and harmonic responses

See Experiment 1.

### Oscillatory coding in fundamental and harmonic responses

We applied multiclass LDA to the epochs of MEG data where participants were viewing the four videos. Every channel and trial was individually baseline-corrected by subtracting the mean of the amplitude between 250ms and 50ms before video onset. To speed up computation, the MEG data were downsampled to 500Hz and re-epoched so they began 250ms before video onset and ended 1,000ms after video onset. Then, for each trial and sample individually, topographic patterns were z-scored across sensors.

The LDA classifier was trained using data from all conditions except one stimulation condition. The classifier was then tested using the held-out stimulation condition. Decision values were computed for every trial, providing an interval measure of classifier performance. This process was repeated 20 times, shuffling the labels of the training dataset to generate a surrogate distribution of the decision values expected by chance.

For time-series analysis of LDA performance, the true decision values of every participant were normalised by subtracting the mean of the surrogate distribution and dividing by the surrogate distribution’s standard deviation. The data were then pooled and subjected to a cluster-based permutation test (1,000 permutations). LDA performance was considered significant when the resulting p-value was less than 0.05.

For the spectral analysis of LDA performance, we focused our analyses on activity between 100ms and 400ms post-stimulus (that is, when the classifier peaked), reasoning that samples outside of this window had little, if any, evidence of representation content. This reduced window led to unreliable EMD derivations, so we used Morlet wavelets to derive the power spectrum. Given that (i) the preceding analyses demonstrate the presence of true harmonic oscillations and (ii) the control conditions provide alternate means to separate fundamental and harmonic responses, we do not believe that this Fourier-based approach undermines the results of this section. We used wavelets at 200 equidistant frequencies between 1Hz and 100Hz, all of which had a length of half of the wavelet’s frequency. As above, this process was repeated for each surrogate, with the true power spectrum being normalised frequency-by-frequency using the mean and standard deviation of the surrogate distribution.

The resulting power spectrum produced a 1/f power law, including positive and negative values. Negative values cannot be easily incorporated into 1/f correction procedures (e.g., FOOOF), so rather than removing the 1/f, we computed the change in power at the target frequencies (34Hz and 68Hz) relative to neighbouring frequencies (i.e., ±5Hz). Decoding performance at the target frequency was directly compared to its neighbours in a permutation-based t-test (1,000 permutations). A peak was considered significant if it was significantly larger than the neighbouring frequencies and the p-value was less than 0.05.

These steps were repeated for two control analyses using bandstop filters. The first control suppressed activity at the fundamental frequency using infinite impulse response filters with sidebands at ±10Hz. The second control suppressed the second harmonic activity, using infinite impulse response filters with sidebands at ±10Hz. All other aspects of the analysis remained the same.

### Experiment 3

We re-analysed an existing, preprocessed dataset of seventeen participants. An outline of the experimental paradigm and stimulation protocol is presented in Figure 2C-D. For further details, see the original paper^83^.

### Participant exclusion

Four participants did not pass the inclusion threshold (as described in the methods for Experiment 1), leaving thirteen participants for further analysis.

### Spectral separation analyses

Analyses matched that of Experiment 1, with two exceptions.

1. We reconstructed a photodiode from EEG triggers and used this to detect peaks in the data.
2. This dataset lacked a baseline condition which involved participants performing the task. Therefore, we compared the stimulation conditions to the eyes-open resting state (see original paper for details).

### Spatial separation analyses

Analyses matched Experiment 1, with the exception that the trials came from the three significant conditions: 4Hz, 6Hz and 8Hz.

### Correlations between fundamental and harmonic responses

See Experiment 1.

### Individual diFerence analyses

Individual alpha frequency (IAF) was computed using eyes-open resting state data. EMD was used to estimate the power spectra of the preprocessed EEG. The approach matched that used for the spectral separation analyses, with the exception that we computed power across all IMFs rather than selecting IMFs at the fundamental/harmonic frequencies – this change circumvented the need to make assumptions about the approximate frequency of the endogenous alpha oscillation. IAF was quantified as the largest peak in the average power spectrum for posterior central channels (Oz, O1, O2, POz, PO3, PO4, Pz, P1, P2).

To compute correlations between IAF frequency and 6Hz fundamental/12Hz harmonic responses, we took the fundamental/harmonic responses used for the spectral separation analyses, averaged these responses across channels, and restricted them to the relevant frequency (that is, the fundamental power spectrum was restricted to power at 6Hz; the harmonic power spectrum was restricted to 12Hz). Pearson’s correlations were conducted across participants, for the fundamental and harmonic responses separately. The correlation was considered significant when the p-value was less than 0.05.

### Analyses linking visual perception to fundamental and harmonic neural responses

We began by modelling perceptual performance as a function of luminance change and its second harmonic. To this end, we generated a time series of 100 samples that covered a single flicker cycle. For every sample, we identified trials with stimuli presented at that flicker phase (+/-0.1π) and computed the mean performance across these trials. This produced a time-course of perceptual performance across the flicker cycle (for example, see Figure 6B).

Two sine waves were generated; one at the fundamental flicker frequency and the other at the second harmonic (the harmonic sine was offset by 0.5π so the peaks of the fundamental and harmonic frequencies aligned). For each participant, these were fitted to the perceptual performance time course in two models. The first model consisted of a constant and the fundamental frequency sine wave only. The second consisted of a constant and both sine waves. Akaike information criterion (AIC) and Bayesian information criterion (BIC) were computed for both models. The changes in AIC/BIC between models were pooled across participants and then subjected to a one-tailed, one-sample t-test. Model improvement was deemed significant when the AIC/BIC was consistently lower than zero across participants and the associated p-value was less than 0.05.

To link this back to the complex waveforms observed in the EEG, for each channel individually, we took the first cycle of the fundamental IMF and the first two cycles of the harmonic IMF (to match the duration to the fundamental IMF) of the peak-locked average. These waveforms were interpolated to match the 100 sample-long time-course of the behavioural data, z-scored, and then used in place of the sine waves in the analysis above. Cluster-based permutation tests (permutations = 1000) were used to assess whether power AIC/BIC was consistently lower than zero across participants (i.e., p < 0.05).

---

**Supplementary Figure 1.**
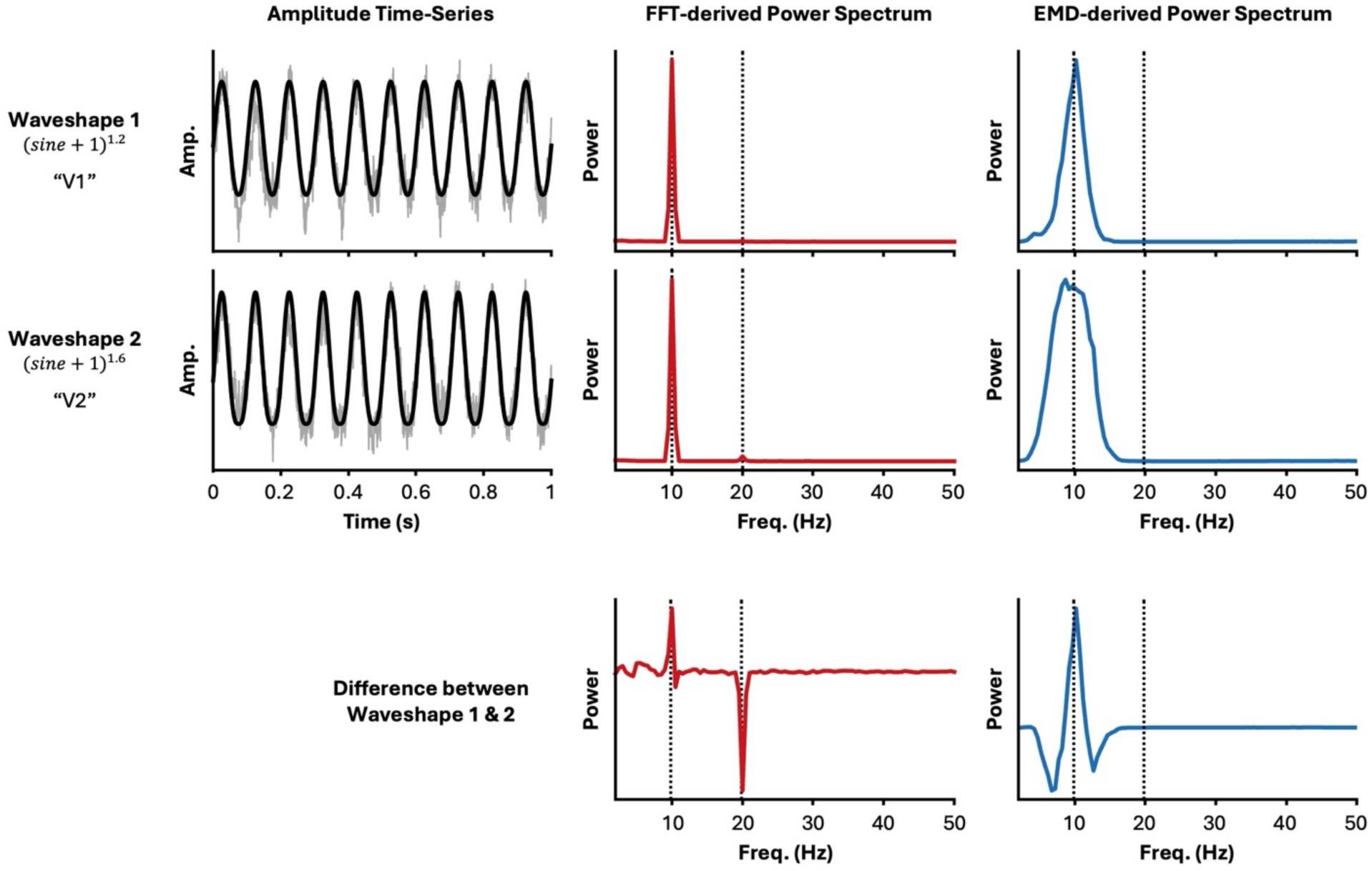
FFT mistakes non-sinusoidal waveforms for multiple oscillations. Past studies have found spatial separation between harmonic components of neural responses to rhythmic stimulation when using the FFT^60–65^. Some may view this as evidence for distinct oscillatory responses to stimulation, but this can simply reflect small changes in waveshape at the fundamental. To demonstrate this, we simulated two non-sinusoidal 10Hz oscillations (left column; black lines) and added pink noise (grey lines) [for the sake of this hypothetical, consider these different waveforms arose in different spatial locations: V1 vs. V2]. The “V2” waveshape deviates more from a pure sine wave than the “V1” waveshape, so it can be considered more non-sinusoidal. We applied FFT (middle column) and EMD (right column) to both. The FFT produced a small but perceptible harmonic at 20Hz for the “V2” waveshape that was driven by the non-sinusoidal waveform. EMD produced no such harmonic. When directly contrasting the power spectra of the two waveforms (bottom row), the FFT showed greater 10Hz power for “V1” and greater 20Hz power for “V2” (mimicking earlier results^60–65^). This suggests that spatial separation methods applied to FFT power spectra could lead to the misinterpretation of differences in waveshape as the presence of a harmonic oscillation. Contrasts of the EMD spectra found no harmonic effect (bottom right), suggesting EMD-based analyses are less susceptible to this problem. Note that while the EMD analyses introduce some quirks of their own (namely that the more non-sinusoidal “V2” waveshape produces a power spectrum with a wider distribution around 10Hz than the “V1” waveshape), this doesn’t impact effects at the harmonic: the key focus of the questions of this manuscript.

**Supplementary Figure 2.**
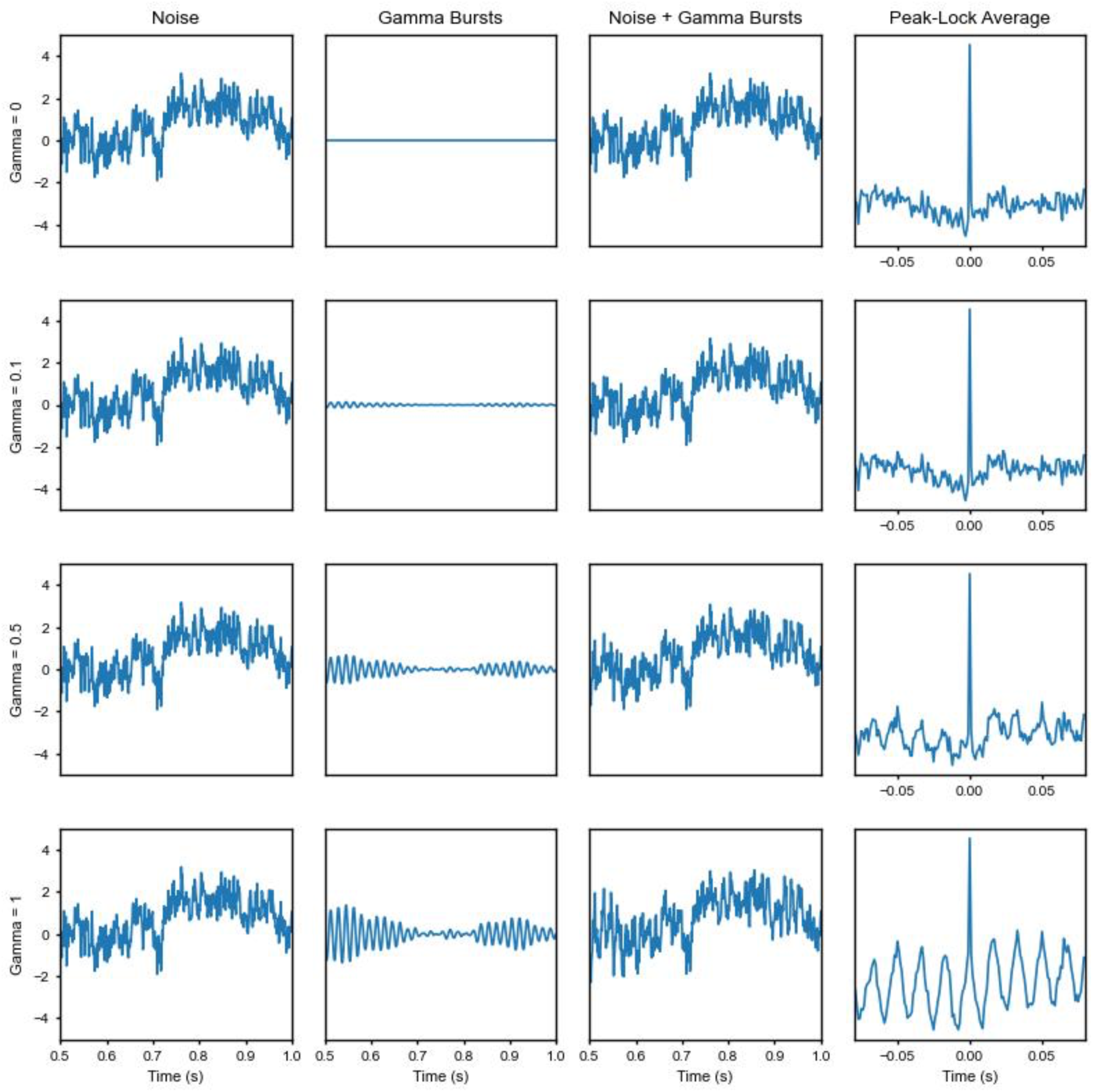
Peak-locking simulation. Low frequencies dominate neural recordings. This makes it difficult to isolate high-frequency responses without resorting to narrowband filtering. Filtering itself, however, can introduce non-sinusoidal rhythmic artifacts, confounding one of the central questions of this manuscript. To circumvent this issue, we use peak-locking: an iterative method that averages the signal across every peak in the data. Some, however, may be concerned that this introduces artificial rhythmicity (that is, rhythmicity where there was none in the original data). To rule out this possibility, we simulated pink noise (far left column) and gamma bursts (middle left column), then summed them together (middle right column). The ratio of the amplitude of gamma bursts relative to the pink noise was varied from 0 (i.e., no rhythmic activity) to 1 (i.e., substantial rhythmic gamma activity). When peak-locking the data (far right column), we found no rhythmicity in the resulting peak-lock average when there was no rhythmic activity (i.e., top row) nor when rhythmicity was very weak (10% of pink noise; second row). This demonstrates that peak-locked averages are not inherently rhythmic and therefore cannot be seen to introduce artificial rhythmicity.

**Supplementary Figure 3.**
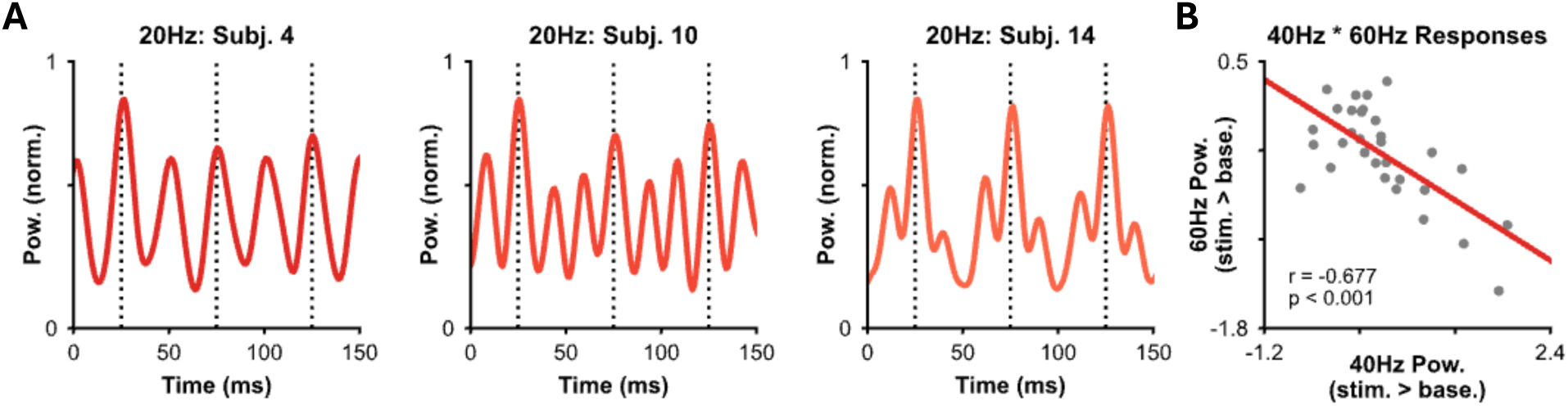
Across-participant variability in harmonic gamma-band responses. **(A)** Peak-lock averages for several participants when receiving 20Hz light stimulation. Some participants show harmonic responses at 40Hz (left; also see Figure 3A), others respond at 60Hz (middle) or display dynamics reminiscent of phase-amplitude coupling (right). **(B)** Participants who show a strong response at 40Hz show weaker responses at 60Hz and vice versa. Dots indicate individual participants. Lines reflect the line-of-best-fit computed using least-squares regression.

**Supplementary Figure 4.**
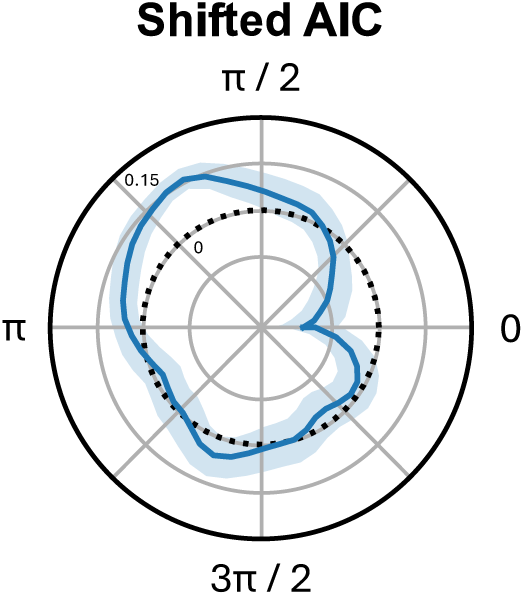
Change in AIC when phase-shifting the harmonic IMF. While AIC does penalise models for increased complexity (theoretically accounting for the difference in regressors between the fundamental and complex models presented in the main text), this penalisation is not always sufficient. Therefore, to complement the analysis presented in the main text, we ran a control analysis where we computed the complex model AIC after phase-shifting the harmonic IMF systematically from zero to 2π radians. If the improvement we saw in model fit for the complex relative to the fundamental model was simply due to the addition of an additional regressor, then AIC should be consistent across all phase shifts of the harmonic IMF. If, however, the observed harmonic IMF is explaining additional variance in the data above and beyond the fundamental model, then the improvement in model fit (i.e., when AIC is lowest) should be greatest for the harmonic IMF without phase-shifting. Indeed, this is exactly what we observed in the polar plot above. In this plot, the dark blue line represents normalised AIC, where values below zero (the black dotted line) indicate a better model fit than the average model fit for all phase shifts. When there is no phase shift (i.e., angle = 0), model performance is significantly greater than the mean AIC across phase shifts (z = –4.77, p = 0.037), indicating that the non-phase-shifted data explains more variability in the behavioural data than would be expected from simply adding any harmonically oscillating regressor. These results support those in the main text and suggest that harmonic alpha activity induced by theta-band stimulation can influence near-threshold visual perception.

